# The pneumococcal two-component system VisRH is linked to enhanced intracellular survival of *Streptococcus pneumoniae* in influenza-infected pneumocytes

**DOI:** 10.1101/767855

**Authors:** Nicolás M. Reinoso-Vizcaíno, Melina B. Cian, Paulo R. Cortes, Nadia B. Olivero, Mirelys Hernandez-Morfa, Germán E. Piñas, Chandan Badapanda, Ankita Rathore, Daniel R. Perez, José Echenique

## Abstract

The virus-bacterial synergism implicated in secondary bacterial infections caused by *Streptococcus pneumoniae* following infection with epidemic or pandemic influenza A virus (IAV) is well documented. However, the molecular mechanisms behind such synergism remain largely ill-defined. In pneumocytes infected with influenza A virus, subsequent infection with *S. pneumoniae* leads to enhanced pneumococcal intracellular survival. The pneumococcal two-component system VisRH appears essential for such enhanced survival. Through comparative transcriptomic analysis between the Δ*visR* and *wt* strains, a list of 179 differentially expressed genes was defined. Among those, the *clpL* protein chaperone gene and the *psaB* Mn^+2^ transporter gene, which are involved in the stress response, are important in enhancing *S. pneumoniae* survival in influenza-infected cells. The Δ*visR,* Δ*clpL* and Δ*psaB* deletion mutants display increased susceptibility to acidic and oxidative stress and no enhancement of intracellular survival in IAV-infected pneumocyte cells. These results suggest that the VisRH two-component system senses IAV-induced stress conditions and controls adaptive responses that allow survival of *S. pneumoniae* in IAV-infected pneumocytes.

**Author summary:** *S. pneumoniae* is an inhabitant of the human nasopharynx that is capable of causing a variety of infections contributing to an estimated 1.6 million deaths each year. Many of these deaths occur as result of secondary *S. pneumoniae* infections following seasonal or pandemic influenza. Although *S. pneumoniae* is considered a typical extracellular pathogen, an intracellular survival mechanism has been more recently recognized as significant in bacterial pathogenesis. The synergistic effects between influenza A and *S. pneumoniae* in secondary bacterial infection are well documented; however, the effects of influenza infections on intracellular survival of *S. pneumoniae* are ill-defined. Here, we provide evidence that influenza infection increases *S. pneumoniae* intracellular survival in pneumocytes. We demonstrate that the poorly understood VisRH signal transduction system in pneumococcus controls the expression of genes involved in the stress response that *S. pneumoniae* needs to increase intracellular survival in influenza A-infected pneumocytes. These findings have important implications for understanding secondary bacterial pathogenesis following influenza and for the treatment of such infections in influenza-stricken patients.

## Introduction

The World Health Organization (WHO) estimates that seasonal influenza virus infections result in about 1 billion infections, 3 to 5 million cases of severe disease, and between 300,000 and 500,000 deaths around the world every year. Oftentimes, influenza infections are complicated by secondary bacterial infections, particularly caused by *S. pneumoniae*. About 11-35% of laboratory-confirmed cases of influenza infection are associated with secondary *S. pneumoniae* infections [1]. Such secondary infections ultimately exacerbate the severity of respiratory symptoms resulting in excess morbidity and mortality [2, 3]. Highlighting the importance of *S. pneumoniae*, it has been proposed that the majority of 40-50 million deaths during the 1918 Spanish influenza pandemic were associated to *S. pneumoniae* secondary bacterial infections [4, 5]. The *S. pneumoniae* is a Gram-positive bacterium of great significance on human health, being the causal agent of otitis, sinusitis, as well as severe diseases such as community-acquired pneumonia, sepsis, and meningitis [6]. More recently, about 34% of the deaths associated with the 2009 pandemic influenza were also linked to secondary bacterial infections, with *S. pneumoniae* as the most commonly associated bacterial pathogen (in addition to *Staphylococcus aureus* and *Streptococcus pyogenes)* [7, 8].

A myriad of concomitant events and factors are thought to be associated with the promotion of secondary bacterial infections following infection with influenza virus: 1) influenza infections produce damage of pulmonary epithelial cells, decreasing the mucocilliary clearance and favoring bacterial adherence and infection [9]; 2) the virus’ neuraminidase results in the desialylation of mucins, which increases pneumococcal adherence [10]; and 3) macrophages and neutrophils infected with influenza virus show impaired phagocytosis of pneumococci [11]. Although these and perhaps other virus-induced modifications on different host cells and tissues [1–3, 8, 12] can contribute to secondary *S. pneumoniae* infections, the precise molecular mechanisms of synergism between influenza viruses and *S. pneumoniae* remain poorly understood.

*S. pneumoniae* is considered a typical extracellular pathogen. However, mounting evidence suggests a significant role of the replication and survival *S. pneumoniae* inside host cells for disease progression and pathogenesis. In this regard, Ercoli *et al* [13] described that intracellular replication of *S. pneumoniae* in splenic macrophages acts as a bacterial reservoir for septicemia. Ogawa *et al* [14] characterized autophagic vesicles that contain pneumococci during the first hours of bacterial infection of human nasopharyngeal epithelial cells and mouse embryonic fibroblasts. The same work also showed that the bacterial protein Ply, a cholesterol-binding, thiol-activated cytolysin, provides advantages for the bacteria to escape from endosomal elimination at early stages of infection. We previously reported that the two-component systems (TCSs) ComDE and CiaRH are involved in the pneumococcal stress response to acidic conditions and in the intracellular survival of *S. pneumoniae* in pneumocytes [15]. In addition, we recently reported that the crosstalk signaling between the serine/threonine kinase StkP and ComE controls H_2_O_2_ production in *S. pneumoniae* modulating its intracellular survival in pneumocytes [16].

In this report, we studied how IAV infection affects the intracellular survival of *S. pneumoniae* in an in vitro pneumocyte IAV-*S. pneumoniae* superinfection model. We observed that *S. pneumoniae* exhibits increased intracellular survival in IAV-infected cells. In *S. pneumoniae*, we identified the two-component system VisRH as a mediator of such increased survival. We found that VisRH controls the expression of 179 pneumococcal genes, such as *clpL* and *psaB*, which encode a molecular chaperone and a Mn^+2^ transporter, respectively. We show that *clpL* and *psaB* expression is required in response to acidic and oxidative stress and for bacterial survival in IAV-infected pneumocytes.

## Results

### Enhanced intracellular survival of *S. pneumoniae* in influenza virus-infected pneumocytes

We previously demonstrated that the *S. pneumoniae* R801 strain can survive inside pneumocytes for several hours [16]. To further define whether a concomitant influenza virus infection would affect *S. pneumoniae* intracellular survival, we established an in vitro IAV-*S. pneumoniae* superinfection model in human-derived A549 pneumocyte cells. As a model virus, we utilized the laboratory-adapted influenza A/Puerto Rico/8/1934 (H1N1) virus (IAV), which has been extensively shown to infect A549 cells [17]. A549 cells were inoculated with a multiplicity of infection (MOI) of either 1, 5, or 10 of the IAV strain. Virus replication was allowed to progress for 24 h before infection with the *S. pneumoniae* R801 strain at a MOI of 30. Flow cytometry using Annexin-V-ACP/PI labeling to test necrosis/apoptosis levels revealed that a MOI of 10 of IAV led to ∼5% increase in the number of necrotic/apoptotic cells compared to non-infected cells (Fig S1A) and ∼15% after bacterial superinfection using a bacterial MOI of 30 (Fig S1B), as described [16]. In further studies, we used IAV at a MOI of 10 and *S. pneumoniae* at a MOI of 30 in the superinfection model. Gentamicin was used to eliminate extracellular bacteria before evaluation of intracellular *S. pneumoniae* following the classical protection assay [16]. Prior IAV inoculation consistently increased bacterial survival by ∼2 fold in A549 pneumocytes (Fig 1A). The synergism between these two pathogens, as defined in this case as enhanced *S. pneumoniae* survival in IAV-infected cells, was also observed in mouse embryonic fibroblasts (MEF) and in cervical cancer cells (HeLa) (Fig 1A), suggesting that this phenomenon is cell-line independent.

**Fig 1.**
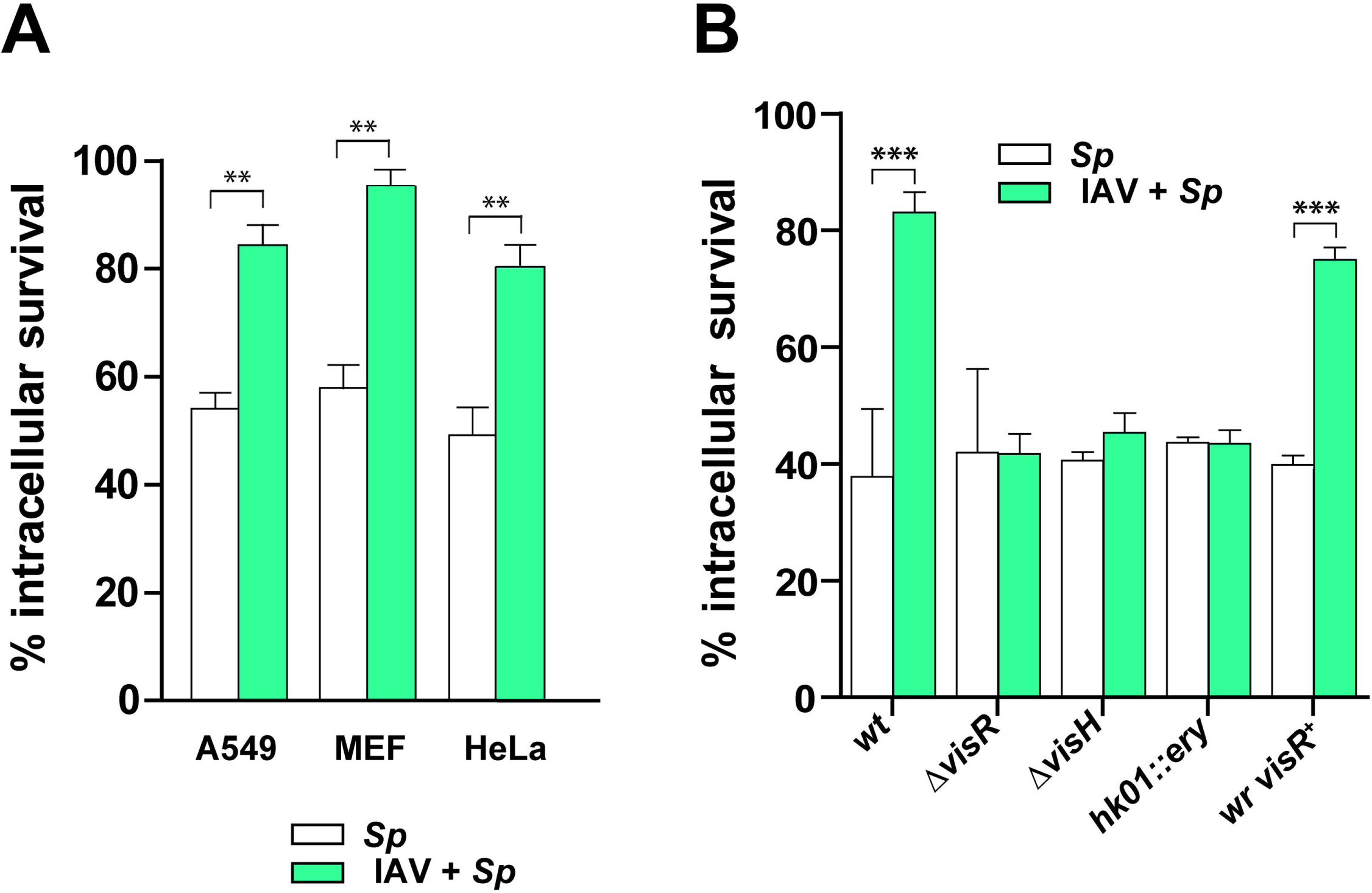
Enhancement of pneumococcal intracellular survival by Influenza A infection is mediated by the VisRH two-component system. (A) The IAV-*S. pneumoniae* synergism is independent of the cell line. The A549, MEF and HeLa cells were treated for 24 h with a viral MOI of 10 and posteriorly infected with the pneumococcal wt strain using a bacterial MOI of 30. Bacterial survival progression was monitored using a typical protection assay. Survival percentages were calculated by considering the total amount of internalized bacteria after 30 min of extracellular antibiotic treatment as representing 100% for each strain. After antibiotic treatment, samples were taken at 4 hours, and pneumocytes were lysed to release pneumococci. Samples were diluted in BHI, spread on BHI-blood-agar plates and incubated at 37°C for 16 h. IAV-infected cells are indicated with green bars, IAV-infected cells with amantadine are indicated with blue bars, and non-virus infected cells with white bars. (B) The synergism between IAV and *S. pneumoniae* is mediated by the VisRH two-component system. A549 cells were previously infected with a viral MOI of 10 for 24 h, and then coinfected by the *wt*, Δ*visH, hk01*::ery (or *visH::ery*) and Δ*visR* strains, and the revertant of the Δ*visR* mutant (*wr visR*^+^). Intracellular survival rates were determined as described in panel A. IAV-infected cells are indicated with green bars and non-virus infected cells with white bars. For all panels, data are representative of at least three independent experiments and statistically significant differences are indicated as *p*<0.05 (*), *p*<0.01 (**) or *p*<0.001 (***).

### The VisRH two-component system mediates enhanced pneumococcal survival in influenza-infected cells

*S. pneumoniae* requires ComE and CiaR response regulators to control the acid stress response and intracellular survival in non-IAV infected A549 pneumocytes [15, 16]. We hypothesized that pneumococcal two-component systems (TCSs) sense physiological changes induced by IAV-infection of pneumocytes and mediate adaptative responses that lead to increased intracellular bacterial survival. Next, we considered that the intracellular changes induced by IAV-infection generate stress conditions sensed by *S. pneumoniae* via TCSs other than ComE and CiaR [15, 16]. From a previous systematic screening of insertion-duplication histidine kinase (*hk*) mutants of *S. pneumoniae* [15] (Table S1), we focused the search on *hk* mutations that were null for pneumococcal intracellular survival in the absence of IAV infection. When non-IAV infected A549 pneumocytes were inoculated with the *S. pneumoniae hk* mutants, most of them showed no changes in intracellular survival compared to the *wt* strain, including the *hk01::ery* mutant (Fig S2). However, in the context of IAV infection, the *hk01::ery* mutant showed impaired pneumococcal intracellular survival compared to the *wt* strain (Fig 1B), indicating that their components participate in sensing the IAV-infected environment. The *hk01::ery* mutant corresponds to TCS01, one of the least studied TCSs but previously identified as a virulent marker in *S. pneumoniae* [18–20]. TCS01, hereafter renamed VisRH (for <u>v</u>irus-<u>i</u>nduced <u>s</u>tress) contains the VisH histidine kinase and the VisR response regulator. Deletion mutants for the *visR* (Δ*visR*) and *visH* (Δ*visH*) genes obtained using the Janus cassette [21] (Table S1), showed similar impairment in intracellular survival as the *hk01::ery* mutant compared to the wt *S. pneumoniae* strain in IAV-infected A549 pneumocytes (Fig 1B). In contrast, the reconstructed revertant of the Δ*visR* mutant (*wr visR*^+^) recovered the *wt* phenotype (Fig 1B). These results confirmed that *S. pneumoniae* needs VisRH for increased intracellular survival in IAV-infected A549 pneumocytes.

### VisRH controls the acidic stress response of *S. pneumoniae*

*S. pneumoniae* needs an acidic stress response for intracellular survival in pneumocytes [15, 16] and survives in acidic autophagic vesicles of Detroit 562 human nasopharyngeal epithelial cells and in mouse embryonic fibroblasts (MEFs) [14]. In IAV-infected cells, *S. pneumoniae* is likely to survive in acidic autophagic vesicles, which implies exposure to the acidic environment and increased ROS production induced by IAV [14]. Since bacterial TCSs typically respond to changes in environmental conditions, we hypothesized that VisRH senses IAV-induced physiological changes at the intracellular level, resulting in an adaptive stress response that improves *S. pneumoniae* survival in autophagic vesicles in IAV-infected pneumocytes [22]. The Δ*visR* mutant was incubated in culture media at pH 4.8 for 1 h showing a 10^3^-fold decrease in bacterial cell viability compared to the *wt.* In contrast, the *wr visR^+^*revertant recovered the acidic tolerance (Fig 2A). The Δ*visR* mutant behaved similarly as the control *atpC^A49T^* mutant, which contains a point mutation at position 49 of the subunit έ of the F_0_.F_1_-ATPase (a proton pump that controls intracellular pH) and is unable to respond to acidic stress in acidified media [15, 23]. These results suggest that VisRH is required for the acidic stress response of *S. pneumoniae*. To further define the role of vesicle acidification in *S. pneumoniae* survival, A549 cells were treated with Bafilomycin A1 (100 nM, 3h), a known v-ATPase inhibitor that halts lysosomal acidification [24] and prevents the fusion between endosome/autophagosome and lysosome, and simultaneously inoculated with *S. pneumoniae*. Intracellular survival of the pneumococcal *wt* strain showed a significant increase when A549 cells were exposed to Bafilomycin A1, as described [25]. In contrast, when Bafilomycin A1-treated or non-treated A549 pneumocytes were infected with either the control *atpC^A49T^* mutant or the Δ*visR* mutant cells (Fig 2B), *S. pneumoniae* showed no increased survival suggesting that the Δ*visR* mutant is unable to respond to the pH variation in vesicles. Since IAV infection also leads to inhibition of the autophagosome/lysosome fusion step [26], VisRH is likely involved in the regulation of stress genes required for pneumococcal adaptation to IAV-induced acidic stress conditions.

**Fig 2.**
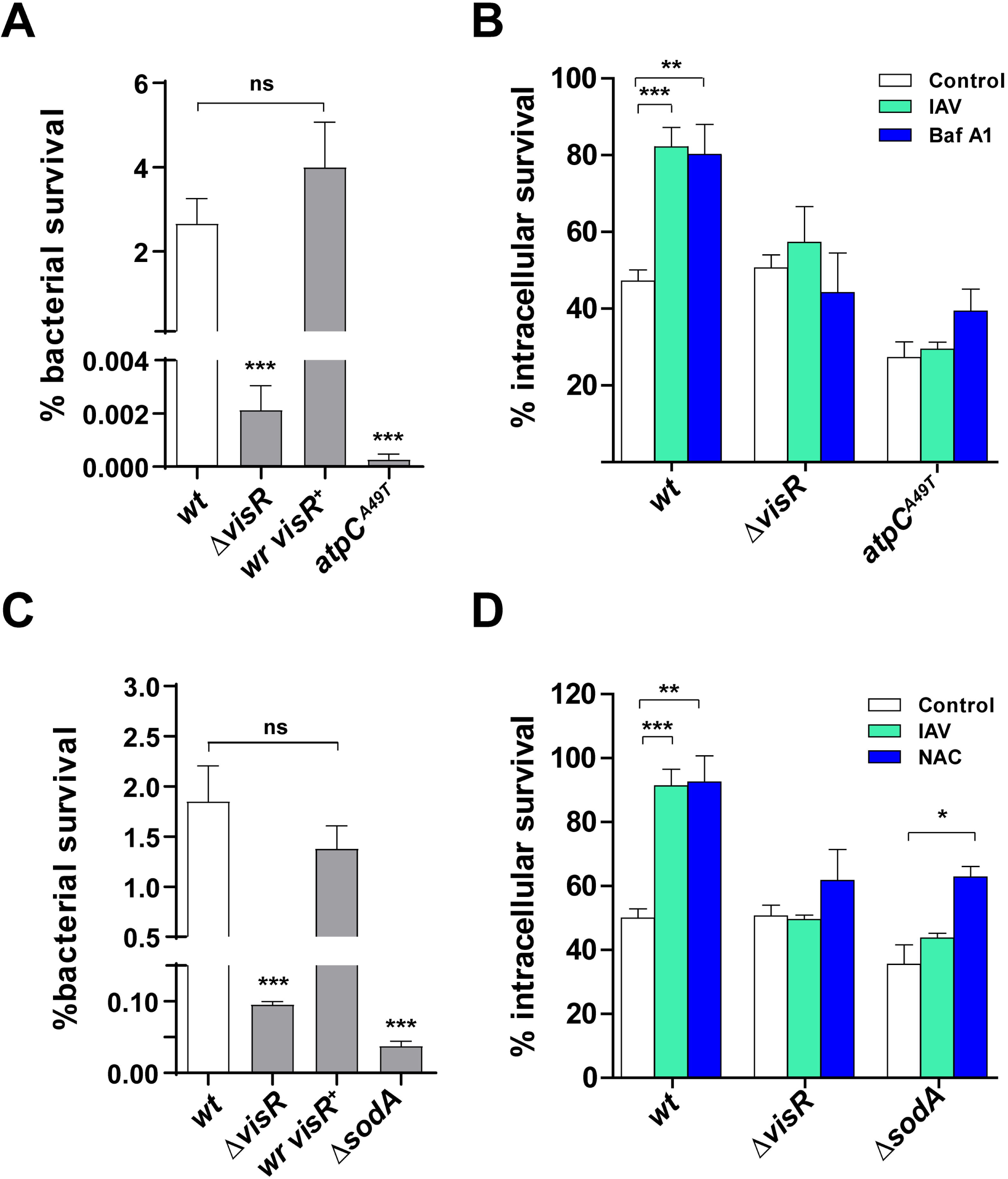
VisRH controls the acidic and oxidative stress response of *S. pneumoniae* in both culture media and pneumocytes. (A) The Δ*visR* mutant is susceptible to acidified media. The Δ*visR*, *wr visR^+^, atpC^A49T^* and *wt* cells were grown in BHI until an OD_620nm_ 0.3 and then incubated in ABM at pH 4.8 for 1 h. Viable cells were assessed by spreading dilutions in BHI-blood-agar plates and incubating these at 37°C for 16 h. (B) Bafilomycin-A1-induced lysosomal neutralization does not affect the impaired intracellular survival of *the* Δ*visR mutant* in IAV-infected cells. A549 cells were infected with the Δ*visR*, *atpC^A49T^* and *wt* cells and intracellular survival was determined as described in the Fig 1. White bars correspond to non-virus infected cells, green bars to IAV-infected cells and blue bars to Bafilomycin-A1-treated cells. (C) The Δ*visR* mutant is sensitive to H_2_O_2_. The Δ*visR*, *wr visR^+^*, Δ*sodA* and *wt* cells were grown in BHI and then exposed at BHI medium containing 20 mM H_2_O_2_ for 2 h. After that, viable cells were determined by spreading dilutions in BHI-blood-agar plates and incubating these at 37°C for 16 h. (D) Inhibition of ROS production does not affect the intracellular survival of *the* Δ*visR mutant*. A549 cells were infected with the Δ*visR*, Δ*sodA* and *wt* cells and intracellular survival was determined as described in the Fig 1 legend. White bars correspond to non-virus infected cells, green bars to IAV-infected cells and blue bars to NAC-treated cells. For all panels, data are representative of at least three independent experiments and statistically significant differences are indicated as *p*<0.05 (*) or *p*<0.001 (***).

### VisRH is involved in the oxidative stress response of *S. pneumoniae*

In a separate study, we previously reported that the StkP/ComE pathway is involved in the regulation of the oxidative stress response that affects the intracellular survival of *S. pneumoniae* in pneumocytes [16]. Additionally, previous reports had indicated that the oxidative stress response is controlled by TCS04 [27], suggesting a complex regulatory system that likely involves the participation of other signal transduction systems. To test the putative role of VisRH in the oxidative stress response of *S. pneumoniae*, we examined the hydrogen peroxide resistance of the Δ*visR* mutant (20 mM H_2_O_2_ in BHI media for 1 h), which was reduced by approximately 30 times while the *wr visR^+^* (revertant) displayed a hydrogen peroxide resistance similar to *wt* (Fig 2C). As a control, we tested the Δ*sodA* mutant (Table S1), a strain deficient in the oxidative stress response that displayed a 10-fold decrease in H_2_O_2_ resistance compared to the *wt* (Fig 2C) [28, 29]. These observations suggest a role of VisRH in the oxidative stress response. IAV-infection of A549 cells leads to enhanced reactive oxygen species (ROS) production and alteration of the antioxidant defense [30, 31]. By measuring the intracellular ROS levels using H_2_DCF-DA, we reproduced this phenotype in our model. We found that ROS production increased by 33% in IAV-infected cells compared to mock-infected cells (Fig S3). In this context, the intracellular survival of the Δ*visR,* Δ*sodA* and *wt* strains was determined in IAV-infected A549 cells. Both the Δ*visR* and Δ*sodA* mutants showed reduced survival rates compared to the wt in IAV-infected cells (Fig 2D), suggesting that VisRH oxidative stress response is relevant for the viral-bacterial synergism. To further explore the effects of ROS production on the intracellular survival mechanism of *S. pneumoniae*, A549 cells were treated with 5 mM N-acetyl-L-cysteine (NAC, 1h prior to S. pneumoniae inoculation), a potent ROS inhibitor [32]. In the absence of IAV infection, the NAC-treated A549 cells lead to increased survival of the *S. pneumoniae wt* strain (∼2-fold) compared to non-NAC-treated A549 cells. In contrast, the Δ*visR* and Δ*sodA* mutants were less sensitive to the effects of low ROS biosynthesis (inhibited by NAC). Overall, VisRH likely senses ROS production to activate an oxidative stress response that allows *S. pneumoniae* to survive into autophagosomes.

### VisR regulates expression of pneumococcal stress genes

Bacterial response regulators control gene expression to develop an adaptive response to stress conditions [22]. In order to identify the VisR-regulated genes, we compared the transcriptomes of the Δ*visR* and *wt* strains by RNAseq analysis. These strains were grown in acidified media at the exponential growth phase, and total RNA was purified and analyzed as described [16]. The transcriptomic analysis revealed the differential expression of 179 genes, 65 were down-regulated and 114 were up-regulated (Fig S4; Table S2; Fig 3A). We identified that VisR controls, directly or indirectly, the expression of stress genes such as those coding for molecular chaperones, redox homeostasis, as well as genes involved in cation and metabolite transport, cell wall biosynthesis, amino acid biosynthesis, purine/pyrimidine, central metabolism, ribosomal and translation structures, among others (Fig. 3B and Table S2). The expression of *visH* showed a 3-fold decrease in the Δ*visR* mutant, suggesting that the VisRH TCS is auto-regulating its own genes (Fig 3C).

**Fig 3.**
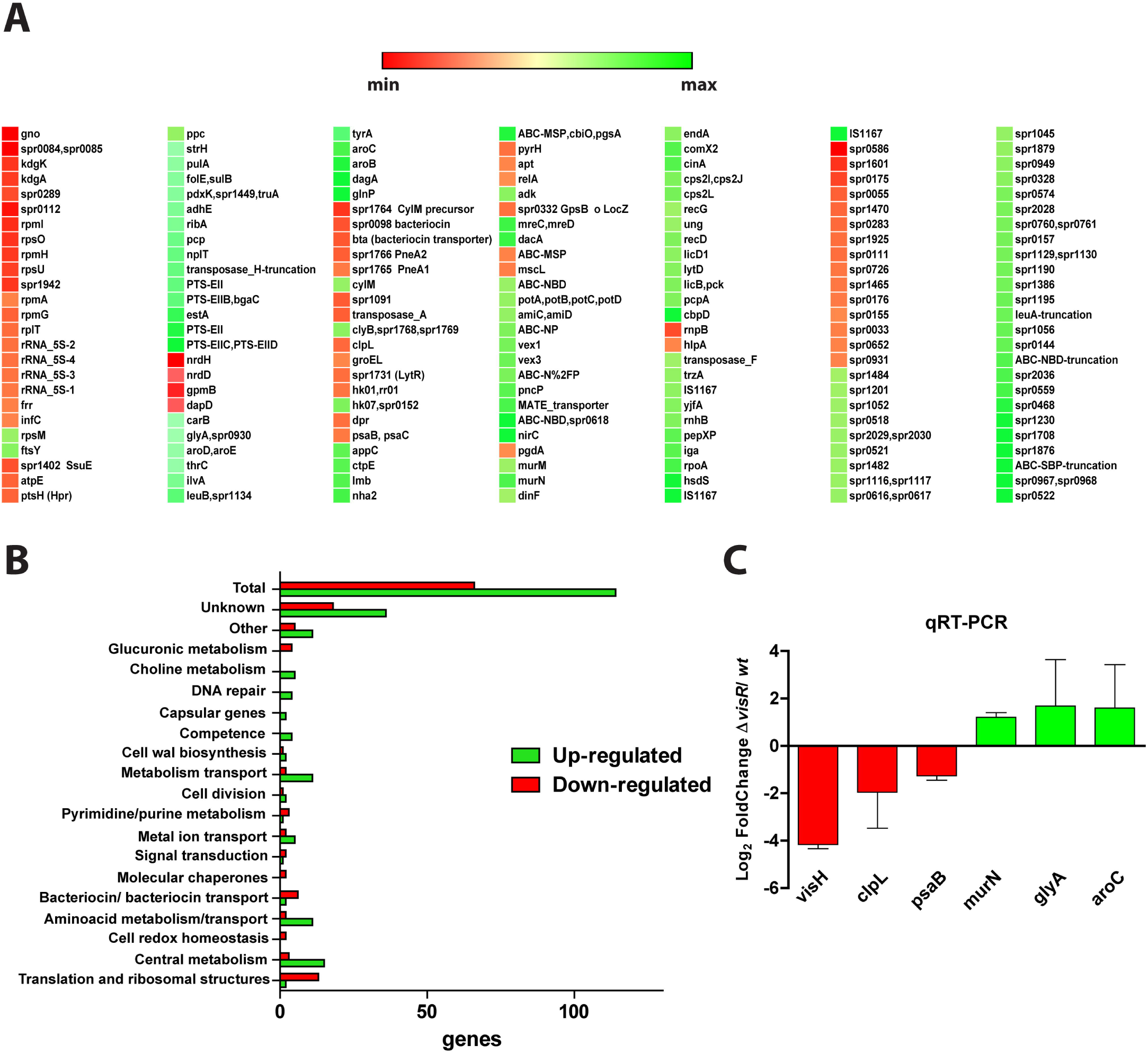
VisR controls gene expression of the stress response in *S. pneumoniae*. (A) RNA-seq heatmap shows gene expression of the comparison between the Δ*visR* and *wt* strains incubated in ABM with relative gene expression in log_2_ fold change demonstrating increased expression in green and decreased expression in red. Gene expressions higher than 2 fold and p values <0.05 were considered significant. (B) Categories of VisR-regulated genes obtained from an RNAseq analysis. An RNAseq generated distribution in functional categories of genes that are regulated in the Δ*visR* mutant relative to strain *wt*. (C) Putative VisR-regulated genes expressed in the Δ*visR* mutant relative to strain *wt.* Gene expression determined by RNAseq was confirmed by qPCR. The Δ*visR* and *wt* strains were grown in BHI to the mid-exponential phase in triplicate and then incubated in ABM for 1h. The fold change in gene expression was measured by RT-qPCR and calculated using the 2^−ΔΔCT^ method. The *gyrA* gene was used as internal control.

We focused on stress genes and confirmed by RT-qPCR that in the Δ*visR* mutant there was decreased expression of *visH* (17.9 times), *clpL* (3.9 times) and *psaB* (2.4 times), and increased transcription of *murN* (2.3 times), *glyA* (3.2 times), and *aroC* (3 times) compared to the *wt* (Fig 3C). The *clpL* gene encodes for a molecular chaperone (heat shock protein) involved in stress response [33, 34], *murN* encodes for an enzyme of cell-wall biosynthesis [35], *glyA* encodes for a glycine hydroxymethyltransferase [36], *psaB* encodes for a subunit of a manganese ABC transporter related to oxidative resistance [27, 29], and *aroC* encodes for chorismate synthase involved in aromatic amino acid biosynthesis in bacteria [37].

To determine a putative correlation between the transcript and the protein levels, we compared the proteomes of the Δ*visR* mutant and *wt*. We used protein extracts obtained from bacterial cells grown in the same conditions described for RNAseq assays. By LC-MS/MS, we detected 925 proteins in total, we found differential expression of 33 down-regulated and 33 up-regulated proteins, and we confirmed the absence of VisR in the Δ*visR* mutant (Fig 4A). The full list of differentially expressed proteins is available (Table S3), and a volcano plot showed proteins differentially expressed with a fold change greater than 2 (Fig S5). When these data were compared with those obtained by RNAseq analysis, we obtained a correlation between the expression of the *clpL*, *psaB*, *dpr* (codes an iron-containing ferritin) [38], *trxA* (encodes for a thioredoxin) [39], *groES* (codes for molecular chaperones) [40], and *nrdD* (codes for a ribonucleotide reductase) [41] genes with the expression of their corresponding encoded proteins (Fig 4B). We found that expression of ClpL and PsaB are repressed in the Δ*visR* mutant, and these proteins have been previously involved in stress response in *S. pneumoniae* [29, 34].

**Fig 4.**
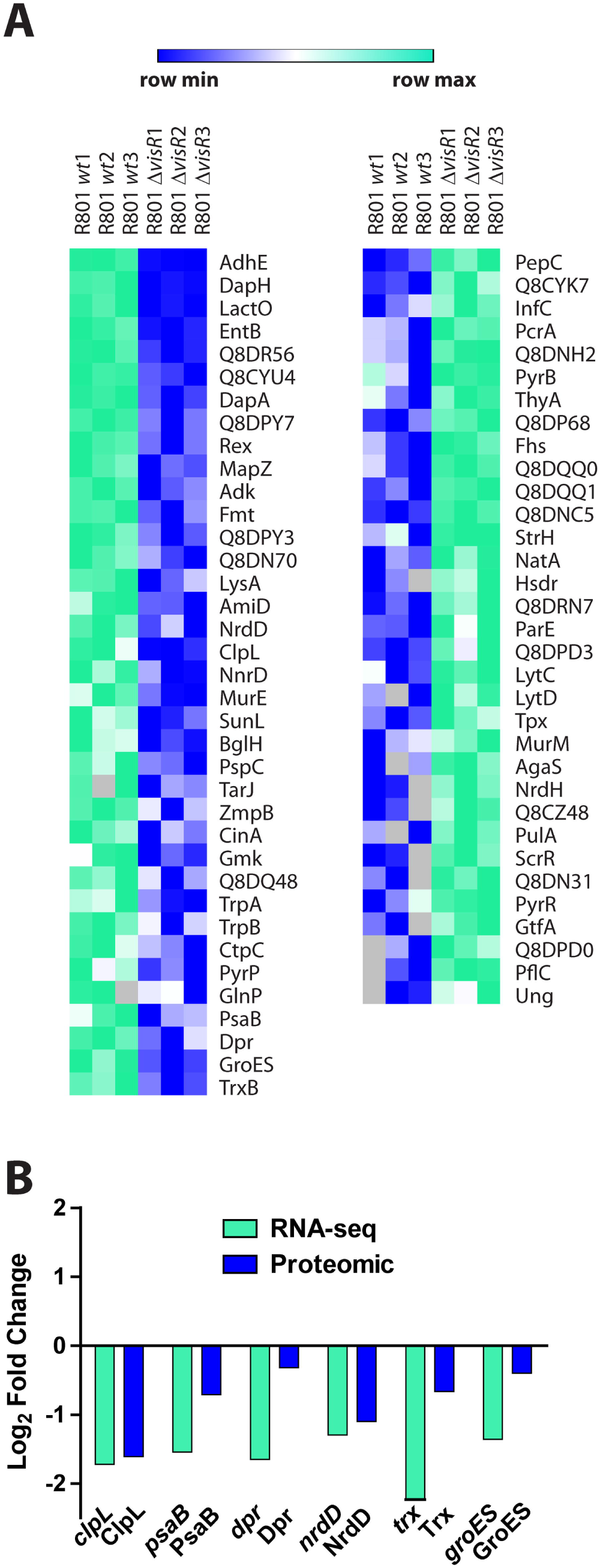
Comparative proteomic analysis of differentially expressed proteins in the Δ*visR* and *wt* strains. (A) Heat map of proteins expressed in the Δ*visR* mutant and referred to *wt*. Proteins with a fold change greater than 2 (less than -1 or greater than 1 on the x-axis of the graph) and a *p*-value < 0.05 were considered as differentially expressed. Higher expression in the *wt* is displayed in shades of green, and higher expression in the Δ*visR* mutant (compared to *wt*) is showed in shades of blue. (B) Comparison between log_2_ folds change (Δ*visR/wt*) obtained by both RNAseq (green bars) and proteomic (blue bars) analysis.

### ClpL and PsaB are involved in the pneumococcal stress response and in the synergistic mechanism between influenza A and *S. pneumoniae*

Since the RNAseq and proteomic data pointed to many stress-related genes under control of VisRH, we focused our study in two particular stress genes, *clpL and psaB.* The *clpL* gene codes for a chaperone that is known to be induced by heat shock [33, 34]. However, experimental evidence showed ClpL is mainly induced under acidic stress (Fig S6). SDS-PAGE comparison of protein extracts from cells grown in ABM (pH 7.8) or incubated in ABM (pH 5.9) showed increased expression of a 78-kDa band under acidic conditions (Fig S6A). Protein sequencing of this band revealed two peptides of 11 and 14 amino acids with 100% homology with the amino acid sequence of ClpL (Fig S6B). ClpL is predicted to have 701 amino acids and a theoretical molecular weight of 77.6 kDa, in line with our observations in SDS-PAGE (78-kDa). To analyze the *clpL* transcript levels under acidic conditions, the *wt* cells were exposed at either pH 5.9 or pH 7.8, and total RNA was purified and treated as described [16]. We detected an increase of 70 times in the *clpL* transcript when cells were exposed to pH 5.9 (Fig 5A) indicating that the rise in ClpL expression is linked to adaptive changes at transcriptional levels that are triggered under acidic conditions.

**Fig 5.**
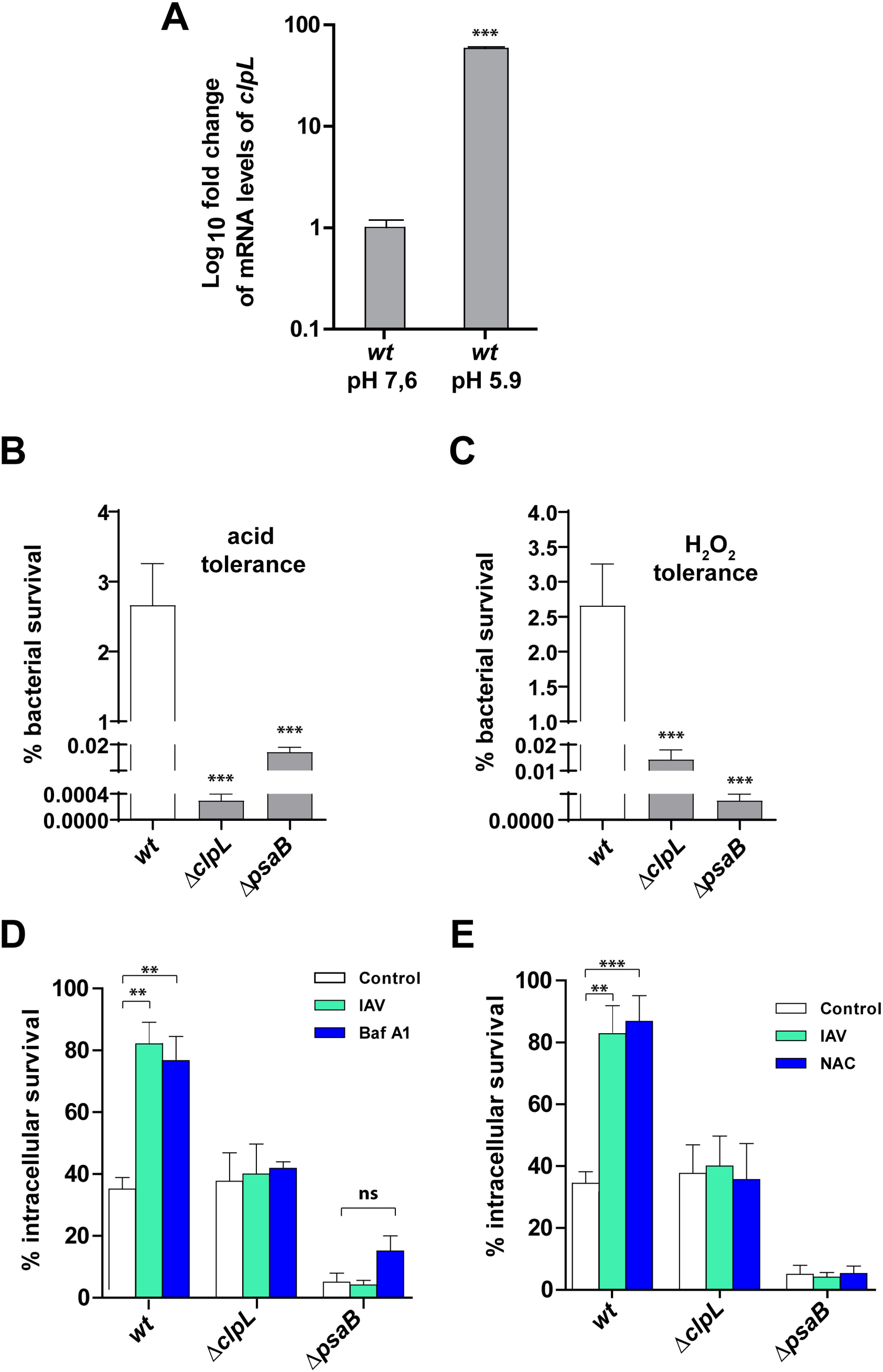
ClpL and PsaB are involved in the pneumococcal stress response needed for the viral-bacterial synergism. (A) Transcription levels of the *clpL* gene increased in cells exposed to acidic pH. The *wt* cells were grown in BHI/pH 7.8 to the mid-exponential phase and resuspended in ABM/pH 5.9, and total RNA was extracted at 1 h. The fold change in gene expression was measured by quantitative real-time PCR and calculated using the 2–ΔΔCT method. The *gyrA* gene was used as the internal control. Error bars indicate the standard deviation of the mean. INSTAT software was used to perform Dunnet’s statistical comparison test for each strain. References: \*\**p*< 0.01; \*\*\**p*< 0.001. (B) The Δ*clpL* and Δ*psaB* mutants are susceptible to acidified media. The Δ*clpL*, Δ*psaB* and *wt* cells were grown in BHI until an OD_620nm_ 0.3 and then incubated in ABM medium at pH 4.8 for 1 h. After that, viable cells were assessed as described in the Fig 2 legend. (C) The intracellular survival of *the* Δ*clpL and psaB mutant* is decreased compared with *wt*. A549 cells were infected with the Δ*clpL*, Δ*psaB* and *wt* cells and intracellular survival was determined as described in the Fig 1 legend. White bars correspond to non-virus infected cells, green bars to IAV-infected cells and blue bars to Bafilomycin-A1-treated cells. (D)The Δ*clpL and* Δ*psaB* mutants are susceptible to H_2_O_2_. The Δ*visR*, Δ*sodA* and *wt* cells were grown in BHI until an OD_620nm_ 0.3 and then exposed at BHI medium containing 20 mM H_2_O_2_ for 2 h. After that, viable cells were determined as described in the Fig 2 legend. (E) Inhibition of ROS production does not affect the impaired intracellular survival of *the* Δ*clpL and* Δ*psaB* mutants. A549 cells were infected with the Δ*visR*, Δ*sodA* and *wt* cells and intracellular survival was determined as described in the Fig 1 legend. White bars correspond to non-virus infected cells, green bars to IAV-infected cells and blue bars to NAC-treated cells. For all panels, data are representative of at least three independent experiments and statistically significant differences are indicated as *p*<0.01 (**) or *p*<0.001 (***).

To define the role of ClpL in the pneumococcal stress response, we constructed a Δ*clpL* mutant (Table S1), which displayed a decrease of 10^4^ times in its tolerance to acidified media (exposure to THYE pH 4.8 for 1 h) compared with the *wt* (Fig 5B). This mutant had the same acid sensitivity as the Δ*visR* mutant, as showed before. With the purpose to determine the effect of oxidative stress, the Δ*clpL* cells were also exposed to H_2_O_2_, which displayed a reduction in H_2_O_2_ susceptibility of 200 times compared with *wt*, indicating that this chaperone is not only a heat shock protein [33] but is also involved both in acidic and oxidative stress responses (Fig 5C).

To determine the contribution of ClpL in our cellular infection model, A549 cells were infected with the Δ*clpL* mutant, and it displayed that its intracellular survival capacity was similar to *wt*. However, when A549 cells were treated with 100 nM Bafilomycin A1 or were previously infected with IAV, conditions that expose pneumococci to the acidic environment of autophagosomes, the Δ*clpL* mutant did not show an increased survival as *wt* did (Fig 5D). Altogether, these findings indicate that ClpL is involved in the acidic stress response, which is in turn required for increased intracellular survival of *S. pneumoniae* in IAV-infected pneumocytes.

Based in the RNAseq assays, we were also interested in the *psaB* gene that encodes for a Mn^+2^ transporter in *S. pneumoniae*. It was reported that the Δ*psaB* mutant displays susceptibility to oxidative stress [29]. We hypothesized that lack of *psaB* could influence the intracellular survival of *S. pneumoniae* in IAV-infected cells due to the virus’ ROS production. The Δ*psaB* mutant strain was 400-fold more sensitive to acidic stress (Fig 5C) and showed 10^4^ times more susceptibility to 20 mM H_2_O_2_, in line with previous studies [28, 29]. In contrast to the Δ*clpL* mutant, Δ*psaB* displayed an impaired intracellular survival in non-IAV infected cells (Fig 5E). However, both mutants failed to exhibit increased intracellular survival in IAV-infected or NAC-treated A549 cells, resembling the phenotype observed for the Δ*visR* mutant (Fig 5E). These observations confirm that ClpL and PsaB are necessary for the IAV-*S. pneumoniae* synergistic mechanism.

### Influenza A-*S. pneumoniae* synergism occurs only in autophagy-proficient cells

Previous reports showed that IAV induces autophagy but blocks the last step of the autophagic process [26] [42]. *S. pneumoniae* [14, 43] also induces autophagy but survives in autophagic vesicles [14]. In order to determine whether autophagy is affected in our IAV-*S. pneumoniae* superinfection model, the potential increased accumulation of LC3-II due to autophagy inhibition was evaluated. As controls of autophagy assays, A549 cells treated with either Bafilomycin A1, a well-known inhibitor of the late phase of autophagy as well [44], or rapamycin, a well-known autophagy inducer [45] showed increased LC3-II levels by Western blot analysis (Fig S7A-B). When A549 cells were infected with either the IAV, the pneumococcal *wt* strain, or superinfected, increased LC3-II levels were observed (Fig S7A-B), consistent with previous results [46]. In a separate study, mKate2-hLC3 vectors [47] were transfected into A549 cells and subsequently infected by either IAV, *S. pneumoniae* or superinfected. Confocal microscopy results indicated that any of these treatments induced remarkably high mKate2-hLC3 punctation in A549 cells (Fig S7C), indicating autophagy induction.

To confirm the functional role of autophagy in this viral-bacterial synergism, IAV-infected mouse embryonic fibroblasts (MEF *atg5-KO*), which are deficient in autophagy [48], were superinfected with the pneumococcal *wt* strain. As control, similar treatments were performed in the parental cell line (MEF *wt*), which are autophagy-proficient cells. A significant increase in the intracellular survival of *S. pneumoniae* in IAV-infected MEF cells was observed similar to that observed in A549 cells (Fig 6A). In contrast, S. *pneumoniae* superinfection of IAV-infected MEF *atg5-KO* cells showed a significant decrease in bacterial intracellular survival compared to the bacterial infection only (Fig 6B). Similarly, the Δ*visR* mutant showed lower intracellular survival in non-IAV infected MEFs *atg5*-KO relative to non-IAV infected MEFs *wt,* although it was equally deficient in both MEFs wt and MEFs *atg5*-KO previously infected with IAV (Fig 6A-B), as observed in IAV-infected A549 cells. Altogether, these results suggest that VisRH mediates the synergistic mechanism between IAV and *S. pneumoniae* and that this phenomenon occurs only in autophagy-proficient cells.

**Fig 6.**
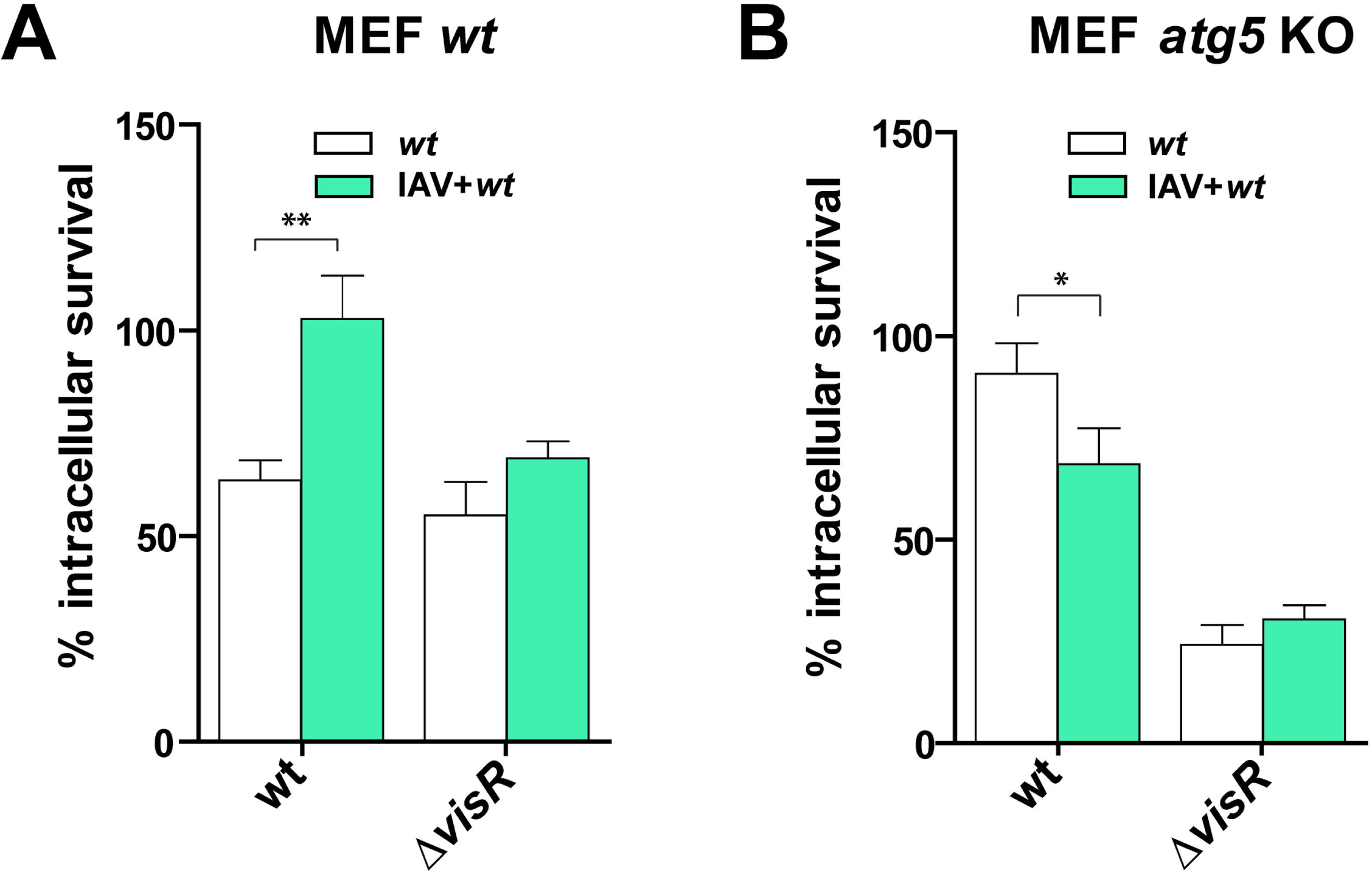
The viral-bacterial synergism is dependent on autophagic-proficient cells. IAV-infected and non-virus MEF (A) and MEF *atg5* KO (B) cells were incubated with the wt and the Δ*visR* strains for 4 h, and bacterial intracellular survival was assessed as described in the Fig. 1A legend. White bars indicate bacterial infection and green bars indicates superinfection. Data are representative of at least three independent experiments and statistically significant differences are indicated as *p*<0.05 (*) or *p*<0.01 (**).

## Discussion

Although *S. pneumoniae* is a common extracellular colonizer of the human nasopharynx, it is known to cause otitis, sinusitis and invasive infections such as pneumonia, bacteremia, and meningitis. Bacterial pneumonia caused *by S. pneumoniae* in patients infected with influenza A has significant relevance in human health during seasonal and pandemic influenza. IAV infections cause physical and physiological changes in the respiratory epithelium that facilitate secondary bacterial infections [10]. Recent reports suggest that such infections are associated with the pneumococcal ability to survive intracellularly. In this regard, Ogawa described intracellular fates of *S. pneumoniae* and found that it is entrapped in specific autophagic vesicles in MEFs [14], which is consistent with our pneumocyte infection model [16]. In the present work, we expanded these studies and found that intracellular pneumococcal survival is clearly improved in IAV-infected pneumocytes.

Many bacterial TCSs, have been involved in intracellular survival mechanisms in eukaryotic cells, such as EvgSA in *Shigella flexneri* [49]; ArcAB [50], PhoPQ [51] and EnvZ/OmpR [52] in *Salmonella thyphimurium*; SrrAB [53]; GraSR [54] in *Staphylococcus aureus*; PhoPQ in *Escherichia coli* [55]; BvrSR in *Brucella abortus* [56]; and PrrAB in *Mycobacterium tuberculosis* [57], among others. In *S. pneumoniae*, most of the TCSs are required for full virulence in animal models of infection [58, 59] and we have shown that two of these systems, StkP/ComE and CiaRH [15, 16], are important in response to acidic and/or oxidative stress. The poorly studied VisRH system (TCS01) has been previously involved in mediating virulence in intranasally-infected mice [18, 19], and a rabbit endocarditis model [20]; however, the effects on intracellular pneumococcal survival were not explored. Here, we show that the intracellular pneumococcal survival of the Δ*visR* mutant is similar to the *wt* in A549 pneumocytes, suggesting that this system may not be directly linked to virulence in animal models.

A key finding of our work was that VisR, as well as ClpL and PsaB, are involved in the stress response induced by *S. pneumoniae* and are necessary for the increased pneumococcal survival in IAV-infected cells. ClpL was formerly described as a heat-shock chaperone induced in pneumococcal cells when incubated at 45°C [34]. The Δ*clpL* mutant is temperature sensitive (to 43°C) but virulence remains unaffected in a murine intraperitoneal model [60]. We observed that ClpL is mostly induced at pH 5.9 in the *wt* strain. In contrast, the Δ*clpL* mutant does tolerate an acidic pH of ≥4.8 in bacterial culture media. A similar phenotype was reported for *Streptococcus mutans* where ClpL was also induced at pH 5.0 and it was essential for the acid tolerance response [61]. We also found that the Δ*clpL* mutant displayed susceptibility to hydrogen peroxide, indicating that ClpL is likely a chaperone involved not only in thermal but also acidic and oxidative stress responses. Previous reports showed that ClpL’s activity is Mn^+2^-dependent [62], further adding to its potential relevance of these proteins in the general stress response of *S. pneumoniae*. Our studies suggest that that ClpL is a key chaperone related to the general stress response of *S. pneumoniae* and essential for bacterial intracellular survival in IAV-infected cells.

PsaB was previously described as an ATP-binding protein that belongs to the ABC-type manganese permease [63]. Mutations on the genes that constitute the *psaBCA* operon result in growth limitations in culture media with low Mn^+2^ concentration, and attenuation in four different animal models of infection [64]. The PsaBCA complex is indeed a Mn^+2^ transporter and its protein components are involved in virulence, resistance to hydrogen peroxide and superoxides [29]. The *psaBC* mutant shows hypersusceptibility to hydrogen peroxide and superoxides [65]. We observed the same phenotype in our Δ*psaB* mutant, confirming that this strain is more susceptible to exogenous hydrogen peroxide than the *wt* strain. Since the *S. pneumoniae* Δ*visR*, Δ*clpL* and Δ*psaB* mutants showed alterations to both acidic and oxidative stress conditions, it suggests a common strategy to general stress adaptation that involves, at least, a TCS, a chaperone and a Mn^+2^ transporter. Such cross-response mechanisms are not unique to *S. pneumoniae*. In *Streptococcus mutants,* a cross-response effect between acidic and oxidative stress was reported for a mutant of the oxidative stress regulator SpxA. Similar to the *S. pneumoniae* Δ*visR* mutant, the *spxA* mutation impairs *S. mutants’* ability to grow under acidic and oxidative conditions [66].

We observed that VisR controls transcription of the *clpL* and *psaB* genes by unknown mechanisms. The VisR response regulator modulates the acidic/oxidative stress response of *S. pneumoniae* to improve intracellular survival in influenza-infected cells. However, transcription of the *clpL* and *psaB* genes could be co-regulated by other regulators [62]. For example, the conserved repressor CtsR regulates the *clpL* expression in many streptococci and lactococci, and these bacteria present CtsR box elements in the *clpL* promoter region [62]. In *S. pneumoniae*, CtsR-binding sites were located upstream from the *clpL* gene [67], however, its regulation has not been yet elucidated. Based on the qPCR assays, we suggest that the *clpL* expression is induced by acidic pH and controlled by VisR, but we cannot discard the possibility that other regulators such as CtsR modulate ClpL stress response in *S. pneumoniae*. Equally complex appears to be the regulation of the *psaB* gene. Expression of the *psaBCA* operon is controlled by the PsaR regulator in a Mn^+2^-dependent manner [68]. In addition, RR04, which belongs to the TCS04, is necessary for the activation of the *psaBCA* locus [27]. Here, we demonstrate that VisR is also essential for the transcriptional activation of the *psaB* gene, adding to complexity of this regulation.

Regarding the increased intracellular survival of *S. pneumoniae* in IAV-infected A549 cells, it is clear that VisR is necessary in IAV-infected cells to but not in non-infected cells. It is likely that IAV infection produces stress conditions in pneumocytes that *S. pneumoniae* overcomes in a VisRH-dependent manner, to improve its capacity to survive intracellularly. Based on the bacterial survival assay in acidified media, where the Δ*visR*, Δ*clpL* and Δ*psaB* cells showed impaired acidic tolerance, we suggest that *S. pneumoniae* needs an adaptive process to survive under acidic conditions, such in acidic vesicles in IAV-infected pneumocytes. Taking into account that ClpL expression is induced at pH 5.9 in the *wt* strain and that the Δ*visR* cells show decreased ClpL and PsaB expression, VisRH is likely sensing acidic stress and modulating adaptation to such condition (Fig 7).

**Fig. 7.**
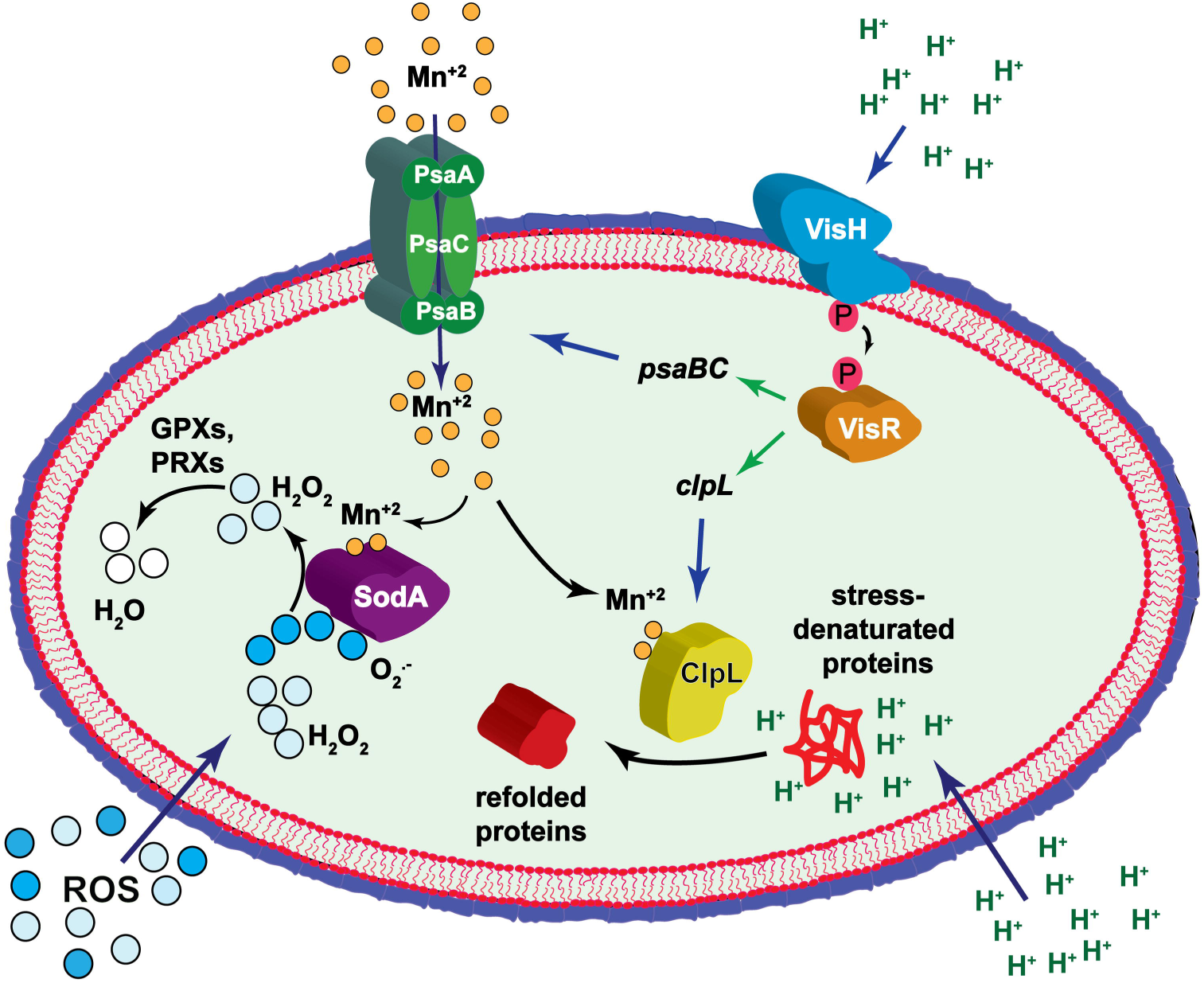
Proposed model for the synergistic mechanism that exists between influenza A and *S. pneumoniae* in pneumocytes.

Related to the putative role of oxidative stress in the synergistic mechanism between IAV and *S. pneumoniae*, it is known that IAV infection increases ROS production in A549 cells [69]. It was reported that *S. pneumoniae* induces an oxidative stress response to survive under oxidative conditions [70]. Analyzing the list of VisR-regulated genes, we focused our attention on the *psaB* gene that encodes for a Mn^+2^ transporter involved in oxidative stress response in *S. pneumoniae* [29, 71]. It was described that the Δ*psaB* mutant was very sensitive to hydrogen peroxide, and this probably occurs due to a low Mn^+2^ level that affects the SodA activity [71]. In view of our results, VisR controls, directly or indirectly, *psaB* transcription affecting the oxidative stress tolerance apparently supported by SodA. To confirm this hypothesis, we tested the Δ*sodA* mutant and we found the same phenotype that the Δ*visR* and Δ*psaB* mutants, which showed increased susceptibility to hydrogen peroxide. Curiously, the Δ*clpL* mutant also displayed an impaired hydrogen peroxide tolerance, suggesting that ClpL is essential for general stress response. Regarding a putative cross stress response, it is important to highlight that ClpL is a Mn^+2^-dependent chaperone [33], in consequence, there is a direct association with the PsaB Mn^+2^ transporter. Probably, in the Δ*psaB* mutant, the observed decreased tolerance to acidic pH that corresponds to diminished activity of ClpL is also due to a low Mn^+2^ level.

The importance of oxidative stress response in the intracellular survival mechanism of *S. pneumoniae* was revealed when ROS production was inhibited in A549 cells by a NAC treatment during bacterial infection. Under these conditions, we clearly observed that the *wt* strain increased its survival, as described for A549 cells [43], indicating that *S. pneumoniae* must overcome this type of stress to survive intracellularly and that this pathogen is susceptible to changes in ROS levels. In the intracellular context of IAV-infected pneumocytes, this virus is able to increase ROS production [31] and, to achieve synergism, *S. pneumoniae* should be also able to overcome oxidative stress.

Our results indicate that the lack of PsaB impaired the intracellular survival of *S. pneumoniae*, even in non-IAV infected cells, probably because Mn^+2^ is needed in for many bacterial processes [72]. In contrast, VisR and ClpL were not essential for this survival mechanism but VisR, ClpL, and PsaB were found to be necessary for the synergism detected in *S. pneumoniae* in IAV-infected A549 cells. We propose that *S. pneumoniae* needs to induce a VisR-controlled adaptive process during superinfection to express chaperones, such as ClpL, to refold proteins denatured by acidic stress and by the IAV-induced ROS production [31], and Mn^+2^ transporter to provide this metal that is essential for chaperone activity, among other cellular processes (Fig 7).

As mentioned before, Gannage *et al* [26] reported that IAV infection produced accumulation of autophagosome by IAV M2-induced blockage of fusion with lysosomes. On the other hand, Ogawa *et al.* [14] reported that *S. pneumoniae* is able to survive in autophagic vesicles. Based on these findings, we hypothesized that the IAV-*S. pneumoniae* synergism may depend on the autophagic process. This putative autophagy-dependence was confirmed using the MEF cell line, with which we reproduced the same synergism found in A549 cells, but not in the MEF *atg5*-KO cells that are deficient in autophagy, indicating that this synergistic mechanism occurs only in autophagy proficient cells.

Pneumococcal pathogenesis has been studied extensively in the last decades. Although *S. pneumoniae* is considered a typical extracellular pathogen, particular attention has been given to the intracellular survival mechanism in the last years, as mentioned before [15, 16] [13, 14]. In this work, we report for the first time that the intracellular survival of *S. pneumoniae* is enhanced in IAV-infected cells, and this synergism occurs in autophagic-proficient cells. For this survival, *S. pneumoniae* needs a physiological adaptation to IAV-induced conditions, and we propose that the VisRH TCS probably senses these changes at intracellular level and controls the expression of ClpL and PsaB, which are needed to tolerate the acidic pH found in intracellular vesicles, as well as the increased ROS level produced by influenza A.

We consider that our results contribute to the knowledge of the intracellular survival mechanism of *S. pneumoniae* in the context of eukaryotic cells infected with influenza A, with a consequent relevance for the management of secondary infections in influenza-infected patients. We propose that intracellular antibiotics should be also considered for the treatment of pneumococcal infections during an epidemic or pandemic influenza A. Many works have described this particular viral-bacterial synergism [1,8–11], and here we provide experimental evidence on how influenza A infections enhance the intracellular survival of *S. pneumoniae*.

## Materials and methods

### Bacterial and viral strains, plasmids, cell lines, and growth conditions

All bacterial strains, oligonucleotides and plasmids used in this study, as well as cloning and mutagenesis procedures, are listed in the supplementary material (Table S1). Oligonucleotide synthesis and DNA sequencing service were performed in Macrogen Inc. (Seoul, South Korea). The growth conditions and stock preparation for the pneumococcal and *Escherichia coli* strains have been reported elsewhere [23], and the transformation assays have been previously described [23, 73]. The influenza virus A/Puerto Rico/8/1934 (H1N1) (IAV) strain was used for superinfection assays. Viruses were grown in embryonated chicken eggs, and the allantoic fluid was collected, aliquoted, titrated in Madin-Darby canine kidney cells (50% tissue culture infective doses [TCID_50_]) and eggs (50% egg infective dose [EID_50_]), and stored at −80°C until used [74].

### Cell lines and culture conditions

The A549 cell line (human lung epithelial carcinoma, pneumocytes type II; ATCC® CCL-185™) was cultured at 37°C, 5% CO_2_ in Dulbecco’s modified Eagle medium (DMEM) with 4.5 g/l of glucose and 10% of heat-inactivated fetal bovine serum (FBS) (Gibco BRL, Gaithersburg, Md.). Fully confluent A549 cells were split once every two or three days via trypsin/EDTA treatment and diluted in fresh media before being cultivated in Filter cap cell flasks of 75 cm^2^ (Greiner Bio-one no. 658175) until passage 6, as described [16]. A549 cells were transfected/co-transfected with pIRES2-EGFP and pIRES2-M2 using JetPRIME® (Polyplus-transfection, Illkirch, France) following the manufacturer’s instructions in serum-free DMEM (Invitrogen) supplemented with 5% of Fetal Bovine Serum (FBS). The MEF (Mouse Embryonic Fibroblast) and the autophagy-deficient MEF *atg5-KO* cell lines were generously provided by Dr. Noboru Mizushima [48]. These cell lines were cultured under the same conditions as described for A549 cells. The mKate2-LC3 plasmids [47] was obtained from Addgene.

### Intracellular survival assays

The intracellular survival assays of pneumococci were performed as reported previously [15, 16] with modifications. Briefly, approximately 1.5 × 10^5^ of eukaryotic cells (A549, MEF, MEF *atg5-KO*, or HeLa cell lines) per well were seeded in 12 well plates and cultured in DMEM (with 5% FBS) and incubated at 37°C for 24 h. Pneumococci were grown in THYE to the mid-log phase (OD_600nm_ 0.3) and resuspended in DMEM (with 5% FBS). Infection of cell monolayers was carried out using a multiplicity of infection (MOI) 30:1. A549 cells were incubated 3 h with pneumococcal strains and cells were washed three times with phosphate-buffered saline (PBS) and it was added fresh DMEM (with 5% FBS) containing 150 µg/ml gentamicin sulfate (US Biological G2030). After a 30 min period, cells were washed three times with PBS. The eukaryotic cells were trypsinized and the occurrence of apoptosis/necrosis caused by pneumococcal infection was quantified by flow cytometry (Annexin V/propidium iodide labeling kit; Invitrogen) giving 5% approximately for all time points analyzed. To determine intracellular survival, cells were lysed by centrifugation for 5 min at 10,000 rpm and the bacterial pellet was resuspended in THYE medium. The number of internalized bacteria at different time points was quantified after serial dilutions and plating on BHI 5% sheep blood agar plates with incubation for 16 h at 37°C. The time scale referred to the time after elimination of the extracellular bacteria by antibiotic treatment. A 100% survival was defined after 30 min of antibiotic treatment (FigS1), and all the samples were referred to this point to calculate the respective percentages.

For intracellular survival determinations in the viral-bacterial superinfection assays, approximately 1.5 × 10^5^ of eukaryotic cells (A549, MEF wt, MEF *atg5-KO,* and Hela cell lines) per well were seeded in 6 well plates, cultured in DMEM (with 5% FBS) and incubated for 24 h. Posteriorly, DMEM was removed from plates, cells were washed three times with PBS and cultured with DMEM containing 1 µg/mL TPCK-treated trypsin for 1 h, and cells were infected with IAV at a viral MOI of 10 at 37°C for 24 h. In parallel, the occurrence of apoptosis/necrosis produced by IAV infection was determined by flow cytometry (Annexin V/propidium iodide labeling kit; Invitrogen) and it was approximately 5%. To perform survival assays in cells previously infected with IAV and treated with amantadine, we carried out the same protocol described above, but we added 50 µM of amantadine (Sigma) at the same time that gentamycin. An analysis was carried out using a confocal laser-scanning microscope (OlympusFV300) with a 100× oil immersion lens, as described [15].

### Susceptibility to acidic and oxidative stress

To determine susceptibility to acidic pH, bacterial cells were grown in Brain Heart Infusion (BHI; pH 7.2) at 37°C until OD_600nm_ ∼ 0.3, centrifuged at 10,000 g for 5 min, resuspended in Todd Hewitt-Yeast Extract (THYE; pH 4.8) and incubated for 1 h at 37°C. To measure susceptibility to oxidative stress, bacterial cells were grown in BHI at 37°C until OD_600nm_ ∼ 0.3, and 20 mM H_2_O_2_ was added to the cultures for 1 h at 37°C. To determine the survival percentage in these assays (acidic and oxidative conditions), serial dilutions were made in THYE (pH 7.8) and plated onto 5% of sheep blood tryptic-soy agar (TSA) plates. After 24 h of incubation at 37°C, colonies were counted to determine the number of survivors. The percentages were calculated by dividing the number of survivors, at pH 4.8 or 20 mM H_2_O_2_, by the number of total cells at time zero before incubation at stressful conditions. Data were expressed as the mean percentage ± standard deviation (SD) of independent experiments performed in triplicate.

### In-gel tryptic digestion and amino acid sequencing of protein bands separated by SDS-PAGE

The protein band of 78 kDa, separated by SDS-PAGE and stained by Coomassie Blue, was cut and the gel slice was incubated in 100 mM ammonium bicarbonate (pH 8.3) containing 45 mM dithiothreitol at 60°C for 30 min. The sample was cooled at RT, and 100 mM iodoacetamide was added followed by incubation at RT in the dark for 30 min. The gel was then washed in 50% acetonitrile-100 mM ammonium bicarbonate with shaking for 1 h, cut in pieces, and transferred to a small plastic tube. Acetonitrile was added to shrink the gel slices and dried in a rotatory evaporator. Then, the gel pieces were treated with 100 mM ammonium bicarbonate (pH 8.3) containing trypsin at a 10:1 ratio (w/w, substrate: enzyme). The sample was incubated at 37 °C for 16 h, and digestion products were extracted twice from the gel with 0.1% trifluoroacetic acid for 20 min. Extractions were loaded into a C18 high-pressure liquid chromatography column (220 × 1 mm), and peptides were eluted with 80% acetonitrile-0.08% trifluoroacetic acid. Selected peaks were applied to a 477A protein-peptide sequencer equipped with a 140 HPLC (Applied Biosystems) and subjected to Edman degradation sequence analysis at the Laboratorio Nacional de Investigacion y Servicios en Péptidos y Proteinas facility (CONICET)[75].

### Mass Spectrometry Analysis

Protein digestion and Mass Spectrometry analysis were performed at the Proteomics Core Facility CEQUIBIEM, at the University of Buenos Aires/ CONICET (National Research Council) as follows. Protein samples were reduced with dithiothreitol (DTT) in 50 mM of ammonium bicarbonate at a final concentration of 10 mM (45 min, 56°C) and alkylated with iodoacetamide in the same solvent at a final concentration of 30 mM (40 min, RT, in darkness). Proteins were digested with trypsin (Promega V5111). After that, the peptides were purified and desalted with ZipTip C18 columns (Millipore). The digests were analyzed by nano-LC-MS/MS in a Q-Exactive Mass Spectrometer (Thermo Scientific) coupled to a nano-HPLC EASY-nLC 1000 (Thermo Scientific). For the LC-MS/MS analysis, approximately 1 μg of peptides were loaded onto the column and eluted for 120 minutes using a reverse phase column (C18, 2 µm, 100A, 50 µm x 150 mm) Easy-Spray Column PepMap RSLC (P/N ES801) suitable for separating protein complexes with a high degree of resolution. The flow rate used for the nano-column was 300 nL min-1 and the solvent range from 7% B (5 min) to 35% (120 min). Solvent A was 0.1% formic acid in water whereas B was 0.1% formic acid in acetonitrile. The injection volume was 2 µL. The MS equipment has a high collision dissociation cell (HCD) for fragmentation and a Q-Exactive Orbitrap analyzer (Thermo Scientific). A voltage of 3.5 kV was used for Electro Spray Ionization (Easy-Spray; Thermo Scientific,). XCalibur 3.0.63 (Thermo Scientific) software was used for data acquisition and equipment configuration that allows peptide identification at the same time of their chromatographic separation. Full-scan mass spectra were acquired in the Orbitrap analyzer. The scanned mass range was 400-1800 m/z, at a resolution of 70000 at 400 m/z and the twelve most intense ions in each cycle were sequentially isolated, fragmented by HCD and measured in the Orbitrap analyzer. Peptides with a charge of +1 or with unassigned charge state were excluded from fragmentation for MS2.

### Analysis of MS data

Q-Exactive raw data was processed using Proteome Discoverer software (version 2.1.1.21 Thermo Scientific) and searched against *Streptococcus pneumoniae* (strain ATCC BAA-255 R6) UP000000586 protein sequences database with trypsin specificity and a maximum of one missed cleavage per peptide. Proteome Discoverer searches were performed with a precursor mass tolerance of 10 ppm and a product ion tolerance to 0.05 Da. Static modifications were set to carbamidomethylation of Cys, and dynamic modifications were set to oxidation of Met and N-terminal acetylation. Protein hits were filtered for high confidence peptide matches with a maximum protein and peptide false discovery rate of 1% calculated by employing a reverse database strategy.

Proteome Discoverer calculates an area for each protein in each condition. To do this it uses the area under the curve of the 3 most intense peptides for a protein. Areas were calculated for each of the three triplicates and normalized. The data obtained for the area for each protein were processed with the Perseus program (Max Planck Institute of Biochemistry, 1.5.5.3 version, available for free) [76] that allows a deeper statistical analysis. Different scatter plots were done according to the compared samples. For each couple of samples, we plotted log *p*-value (-log Student T-test *p*-value A_B) on the y-axis versus Student T-test Difference A_B in the x-axis. Proteins that appear in the volcano plot with a fold change greater than 2 (less than -1 or greater than 1 on the x-axis of the graph) and a *p*-value < 0.05 (above 1.3 on the y-axis of the graph) were considered as differentially expressed.

### RNAseq analysis

Cells were initially grown in THYE medium at pH 7.8 until OD_600nm_ ∼0.3 (log phase), centrifuged at 14,000 g for 10 min at 4°C, resuspended in the same volume in ABM at pH 5.9 [23] and incubated a 37°C for 1h. Then, cells were centrifuged at 14,000 x g for 10 min at 4°C, resuspended in a 1/10 vol of lysis buffer (DOC 1% in 0.9% Na Cl) and incubated 3 min a 37°C until complete lysis. Total RNA from three biological replicates for *wt* and the Δ*visR* mutant were purified by TRIzol reagent according to the manufacturer’s instructions (Fisher Scientific). The RNA for RNAseq assays was obtained as described [16]. Data analysis was performed as reported [62].

### Differential gene expression

The aligned reads were assembled by Cufflinks (version-2.2.1), and then the differentially expressed genes were detected and quantified by Cuffdiff, which is included in the Cufflinks package, using a rigorous sophisticated statistical analysis. The expression of the genes was calculated in terms of FPKM (fragment per kilobase per million mapped reads). Differential gene expression analysis was carried out between *wt* and the Δ*visR* samples.

### Protein analysis by western blots

The A549 cells were lysed and protease inhibitor cocktail added to obtain the whole protein to be quantified. The lysates with protein loading buffer were boiled for 5 min. The supernatants were collected and 40 μg of each sample were loaded onto 15% SDS-PAGE gels and electrophoresed for protein resolution at RT using Tris-Glycine-SDS running buffer at a constant electric field of 100 V cm-1. Posteriorly, proteins were electroblotted onto PVDF membranes, which were blocked for 2 h at room temperature and incubated overnight at 4°C with primary antibodies diluted at 1:1,000 in PBS with 5% bovine serum albumin buffer. After washing 3 times with Tris-buffered saline (TBS) with 0.5% (v/v) Tween, the membranes were incubated for 2 h at room temperature with Alexa-conjugated secondary antibody (1:1,000 dilution) to detect LC3-II and p62. The membranes were imaged under fluorescence mode in an Oddisey CLx Imaging System (LI-COR), and bands were quantified with Image Studio software (LI-COR). Rabbit monoclonal antibody against LC3A/B (D3U4C) XP(R) (12741P) was obtained from Cell Signaling Technology. Rapamycin (R8781; Rapa, mTOR inhibitor), Bafilomycin A 1 from *Streptomyces* (B1793) and Mouse monoclonal anti-beta-actin antibody (A2228) were obtained from Sigma Life Science. Mouse monoclonal antibody against Influenza A M2 protein [14C2] (ab5416) was obtained from Abcam. Recombinant Rabbit monoclonal antibody against SQSTM1/p62 (701510) was purchased to Invitrogen.

### qRT-PCR

cDNA was synthesized from 2 μg RNA using the ProtoScript II First Strand cDNA Synthesis Kit (NEB) following the manufacturer’s protocol, and cDNA was cleaned using the QIAquick PCR Purification Kit (Qiagen). Genes were amplified using the oligonucleotides listed in the Table S4 and PowerUp SYBR Green Master Mix (Applied Biosystem) following the manufacturer’s protocol. Expression was determined relative to AU0158 normalized by *gyrA* (*spr1099*) expression using the ΔΔCt method [77]. The *gyrA* had a similar expression by RNA-Seq for *wt* and the Δ*visR* mutant, and this had been used to normalize the expression in *S. pneumoniae* in other studies [78].

### ROS detection

To asses ROS production, we used 2′,7′-Dichlorodihydrofluorescein diacetate dye (H_2_DCF-DA; Molecular Probes) following the manufacturer’s instructions. Briefly, we infected A549 cells with IAV at MOI 10 as it was indicated above, 24 h post-infection the cells were trypsinized and washed twice with PBS, resuspended with PBS containing H_2_DCF-DA (10 =M) and incubated for 30 min at 37°C. Then, cells were washed and resuspended with PBS and we measured the intensity of fluorescence of the DCF by cytometry.

### Accession numbers

The RNA-Seq data generated from this study are deposited at the NCBI SRA under the accession numbers SAMN08473835 (*wt* strain) and SAMN08473837 (Δ*visR* strain). This data corresponds to the Bioproject PRJNA433281, and the SRA IDs are SRR6679010 and SRR6679012.

## Acknowledgments

We thank Gabriela Furlan, Laura Gatica, Paula Abadie, Pilar Crespo, Alejandra Romero (CIBICI-CONICET) and Carlos Mas (CIQUIBIC-CONICET) for their skillful technical assistance. We thank Dr. Noboru Mizushima (Department of Biochemistry and Molecular Biology, The University of Tokyo, Japan) for the kind gift of MEF *atg5-KO* and MEF *wt*. We thank Alex Saka for technical discussions.

## Supporting Information

**S1 fig. Determination of apoptosis and necrosis levels in A549 cells infected with IAV and/or S. *pneumoniae*.** (A) A549 cells were infected with different MOI of IAV for 24 h and coinfected with a bacterial MOI of 30. Apoptosis/necrosis was measured at the single-cell level by labeling cells with annexin-V-APC and counterstaining with propidium iodide (PI). Representative data are shown and percentage of cells are indicated in each quadrant (lower left: APC^−^/PI^−^, intact cells; lower right: APC^+^/PI, apoptotic cells; upper left: APC^−^/PI^+^, necrotic cells; upper right: APC^+^/PI^+^, late apoptotic or necrotic cells). (B) The bar chart describes the percentual distribution of necrotic, apoptotic and viable cells after infection with different MOI of IAV or with superinfection with *S. pneumoniae*.

**S2 fig. Identification of histidine kinase (*hk*) mutants of *S. pneumoniae* displaying normal intracellular survival in pneumocytes.** A549 cells were infected with different *hk* mutants and its intracellular survival capacity was determined as described for non-virus infected cells in the Fig 1 legend, and these results were compared with those obtained for the *wt* strain. Green bars and blue bars correspond to 2 h and 4 h of incubation after antibiotic treatment, respectively.

**S3 fig. Confirmation of the IAV-induced ROS production in A549 cells.** (A) Representative flow cytometry histogram showing results of H_2_DCF-DA staining (a measurement of ROS levels) of IAV-infected A549 cells or mock-A549 cells. (B) Bar graph depicting results of IAV-infected A549 cells compared with non-infected cells. Data are representative of at least three independent experiments.

**S4 fig. VisR is a global regulator that controls gene expression during the stress response.** (A) Gene expression scatter plot in samples obtained from the *wt* strain and the Δ*visR* mutants, with the *x*-axis representing the gene expression values for the control condition (*wt*) and the *y*-axis representing those for the treated condition (Δ*visR*). Each black dot represents a significant single transcript, with the vertical position of each gene representing its expression level in the experimental conditions and the horizontal one representing its control strength. Thus, genes that fall above the diagonal are over-expressed whereas genes that fall below the diagonal are under-expressed as compared to their median expression levels in the experimental groups. (B) Volcano plot of gene expression in *wt* vs Δ*visR* samples measured by RNAseq. The *y*-axis represents the mean expression value of the log_10_ (*p*-value), while the *x*-axis displays the log_2_ fold change value. Black dots represent genes with an expression 2-fold higher in the Δ*visR* mutant relative to strain *wt* with a *p*-value < 0.05, with red dots signifying genes with an expression 2-fold lower in the Δ*visR* mutant, which are relative to strain *wt* with a *p* < 0.05.

**S5 fig. Comparative proteomic analysis between the *wt* and the** Δ***visR* strains.** Volcano plot reflecting the results from the statistical analysis of differentially expressed proteins quantified among the proteome of the *wt* strain and the Δ*visR* mutant. Statistical analysis was performed by Student t-test and statistical significance was considered for *p*-values < 0.05. Significant values are represented as red dots.

**S6 fig. Identification of the pneumococcal 78-kDa ClpL chaperone expressed under acidic conditions.** (A) SDS-PAGE analysis of protein extracts obtained from the *wt* cells grew at slightly alkaline (pH 7.8) or acidic (pH 5.9) culture media. The protein band subjected to N-terminal sequencing is indicated by an arrow. (B) The N-terminal sequence obtained by Edman degradation were analyzed by tryptic digestion and HLPC-protein sequencer. The m/z values of ions matching peptides derived from the 78-kDa protein band are indicated by numbers. The amino acids sequences corresponding to pick 8 (11 amino acids) and pick 12 (14 amino acids) corresponded to the ClpL chaperone, according to the R6 pneumococcal genome (https://www.uniprot.org/proteomes/UP000000586).

**S7 fig. IAV and *S. pneumoniae* infection and superinfection induce autophagy in A549 cells.** (A) LC3-II levels are induced by superinfection with IAV and S. pneumoniae. A549 cells were infected with IAV (MOI 10), *S. pneumoniae* (MOI 30) and coinfected as described in the Fig.1A legend. As controls, A549 cells were also treated with inducers (rapamycin) and inhibitors (bafilomycin A1) of the autophagy process. Cell lysates were subjected to Western blot analysis using anti-LC3-II, anti-beta-actin antibodies with data being representative of at least three independent experiments. (B) Quantification of the LC3-II level in western blot: bar graphs represents LC3-II relative intensity (LC3-II/β-actin) with data being representative of at least three independent experiments. (B) Formation of the puncta of mKate2-hLC3 indicating autophagy induction during IAV, *S. pneumoniae* and superinfection. The A549 cells were transfected with the mKate2-hLC3 plasmids for 24 hours, and followed by an IAV, *S. pneumoniae* or superinfection for 4 h. The far-red (mKate2) fluorescence in the cells were monitored using an Olympus FluoView FV1000 confocal laser scanning microscope.

