## Supplementary figures and images for "The pneumococcal two-component system VisRH is linked to enhanced intracellular survival of *Streptococcus pneumoniae* in influenza-infected pneumocytes"

### Fig S1

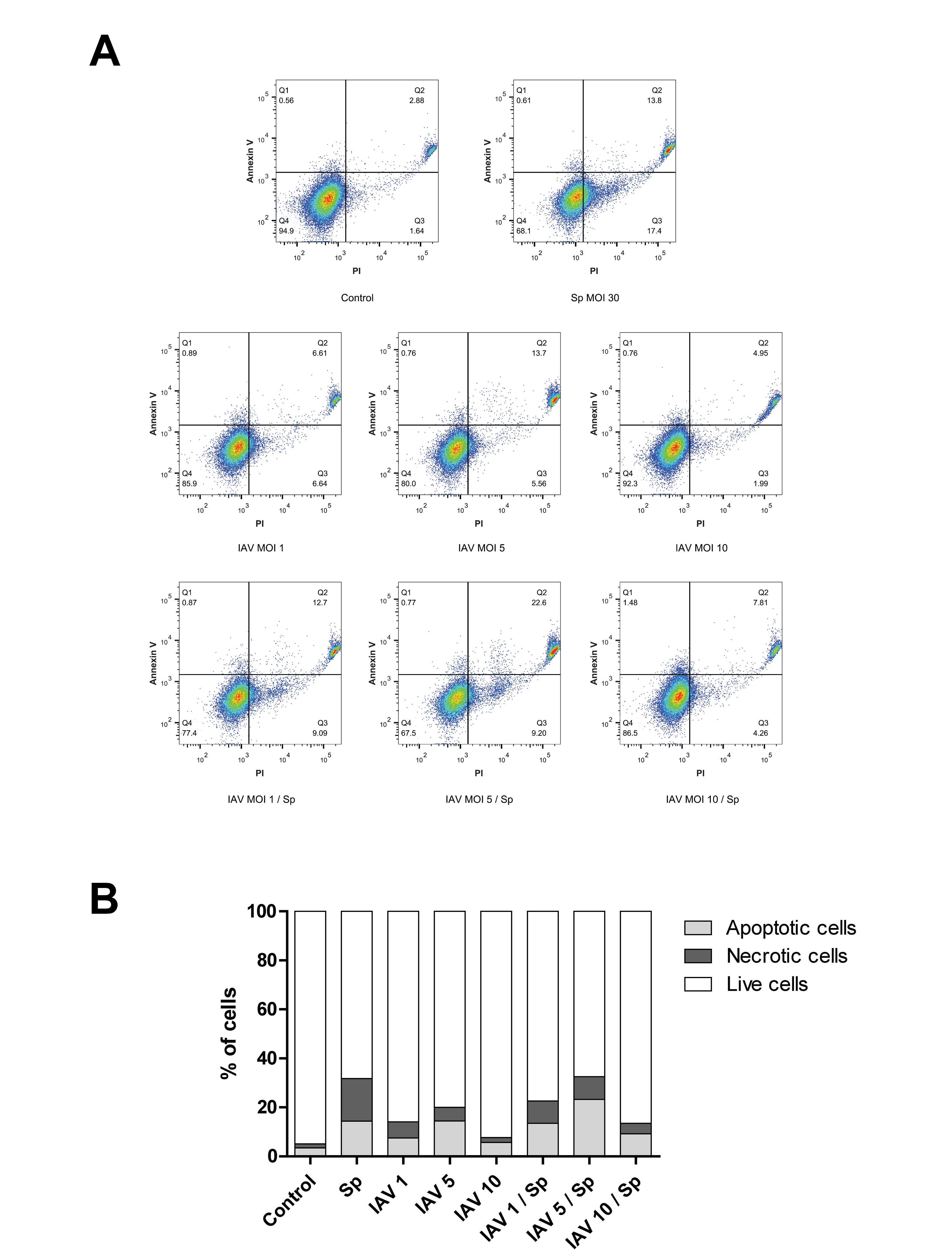

### Fig S2

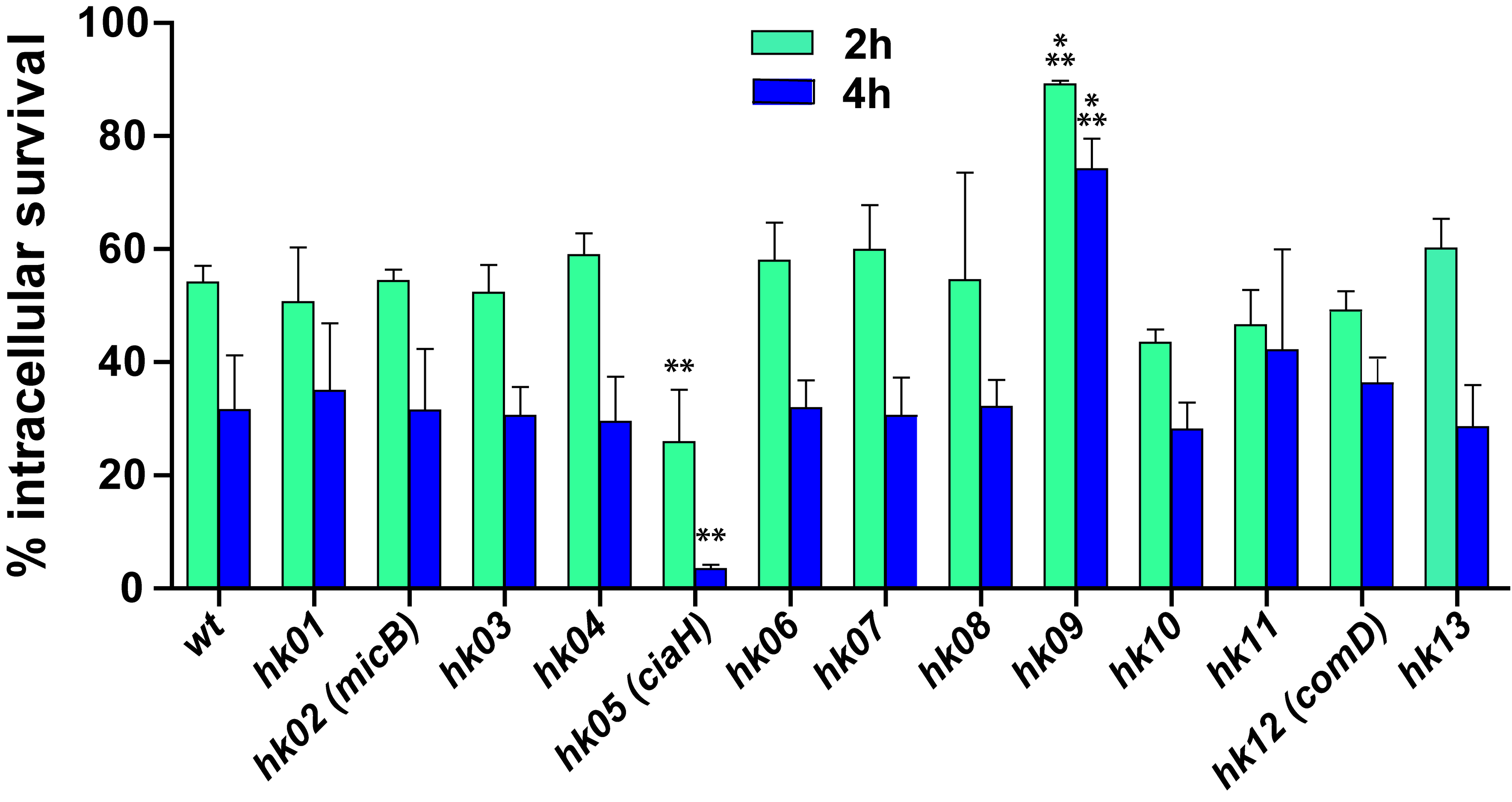

### Fig S3

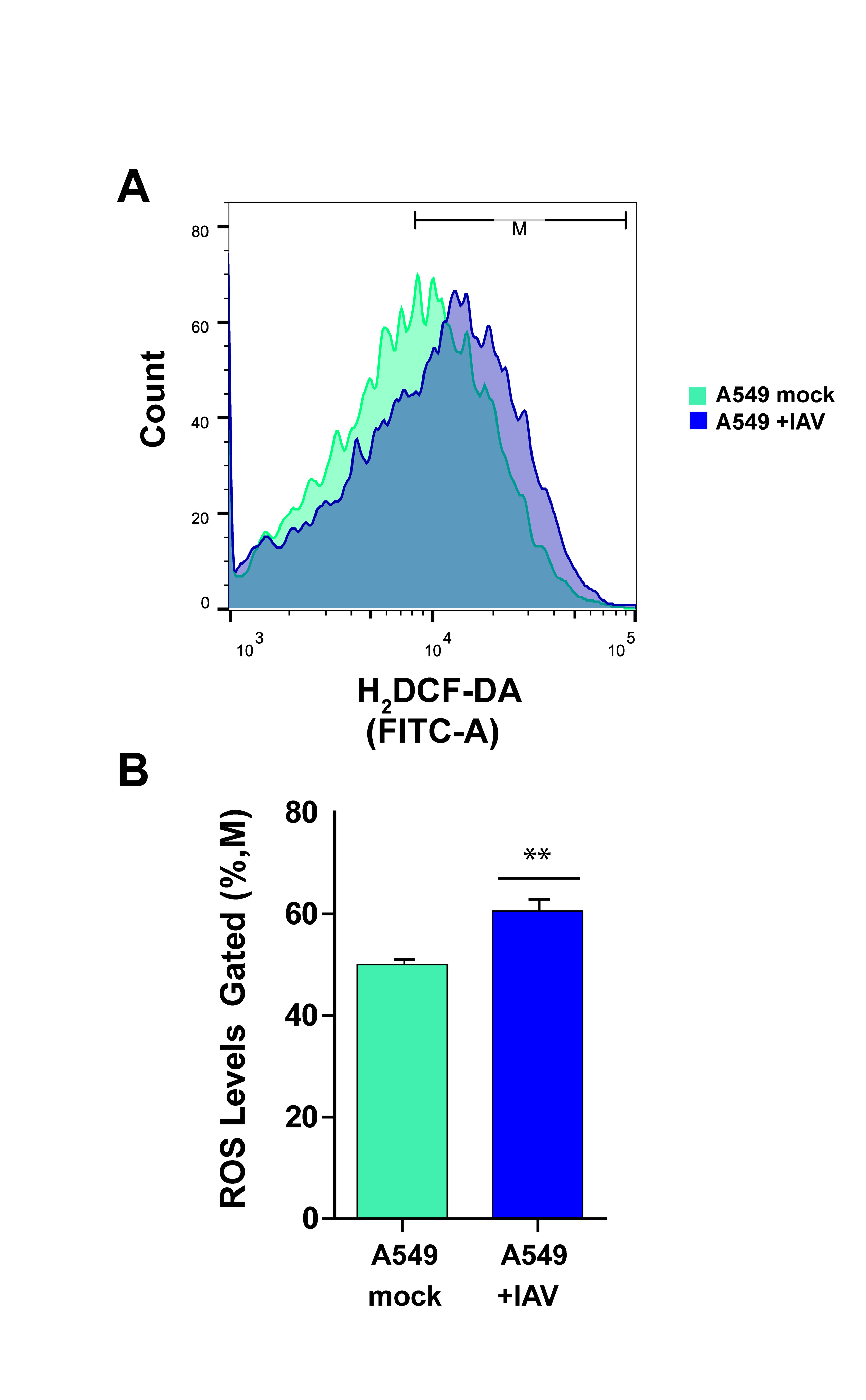

### Fig S4

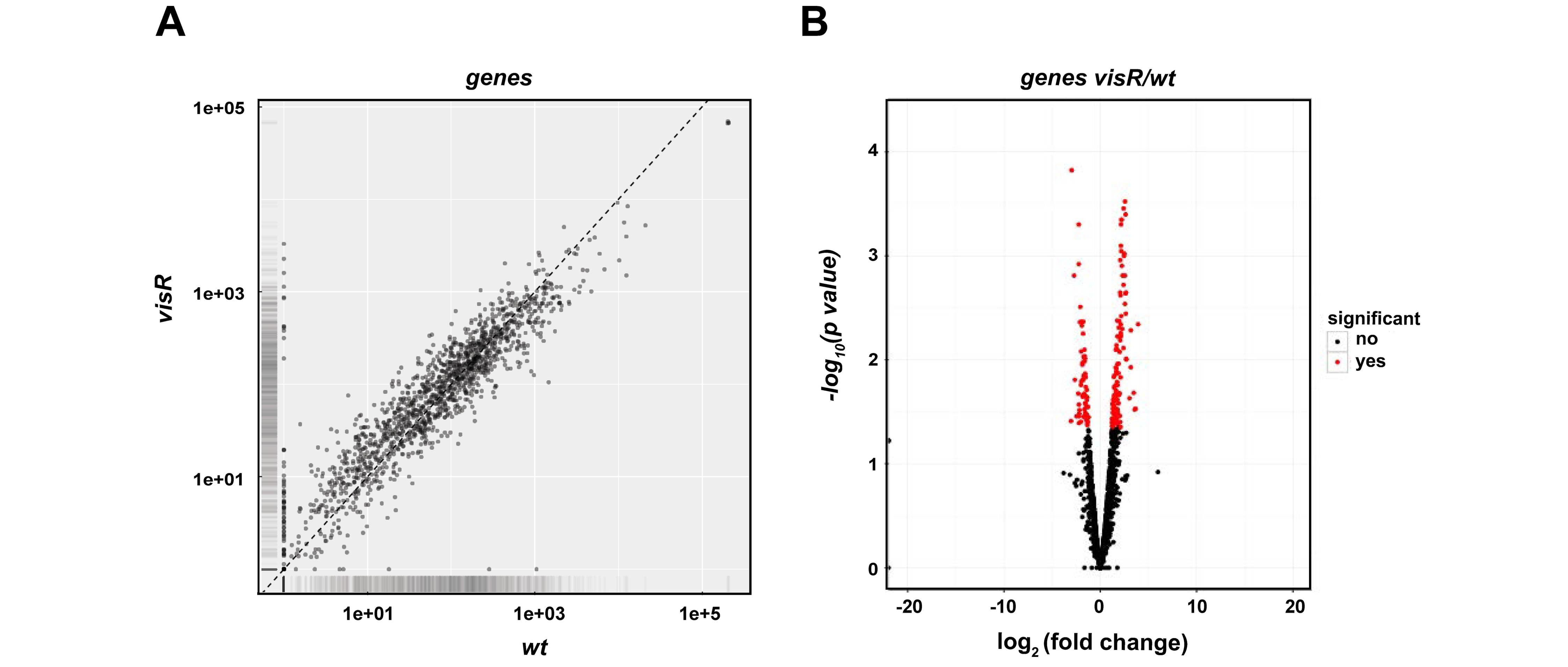

### Fig S5

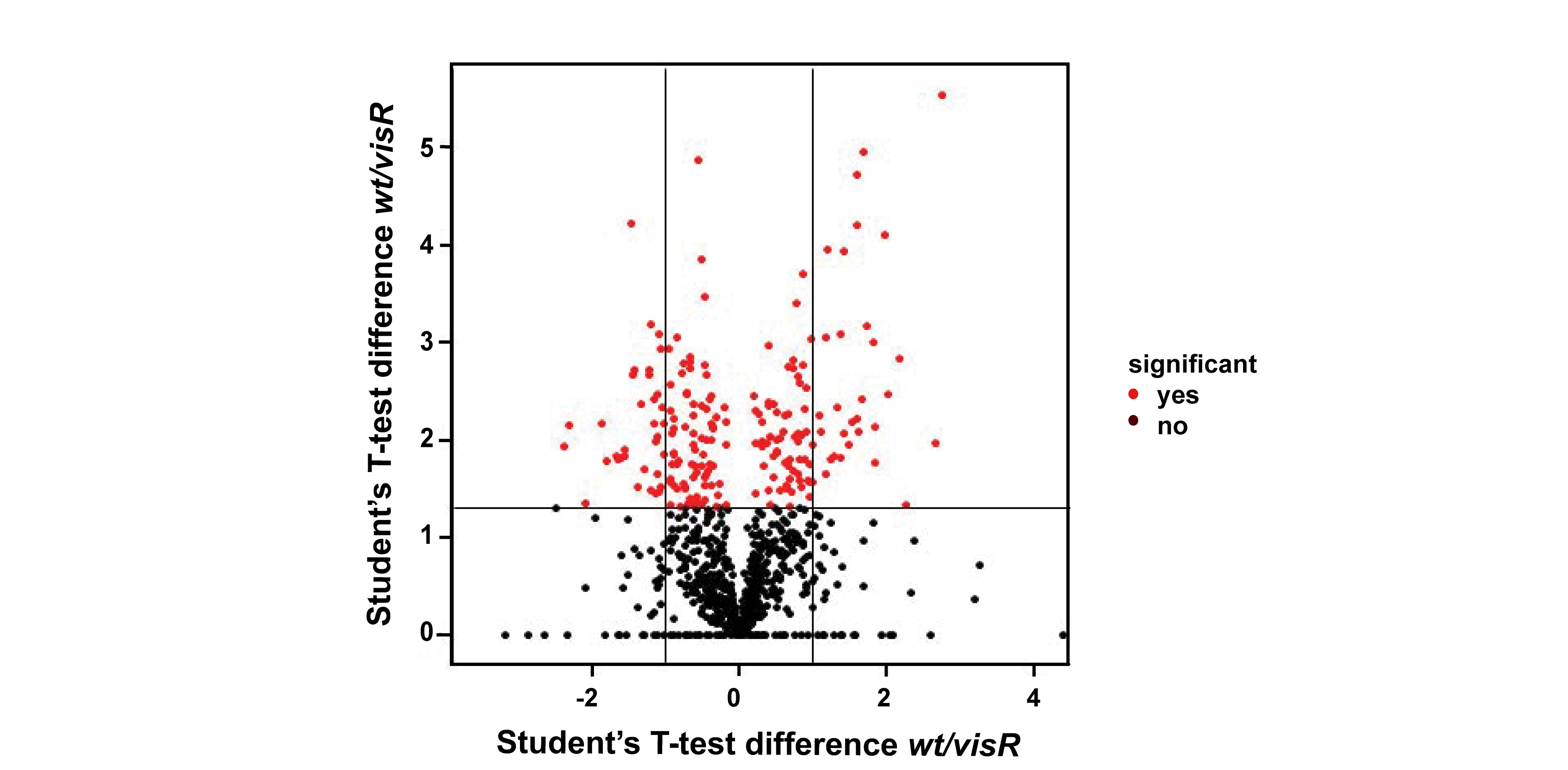

### Fig S6

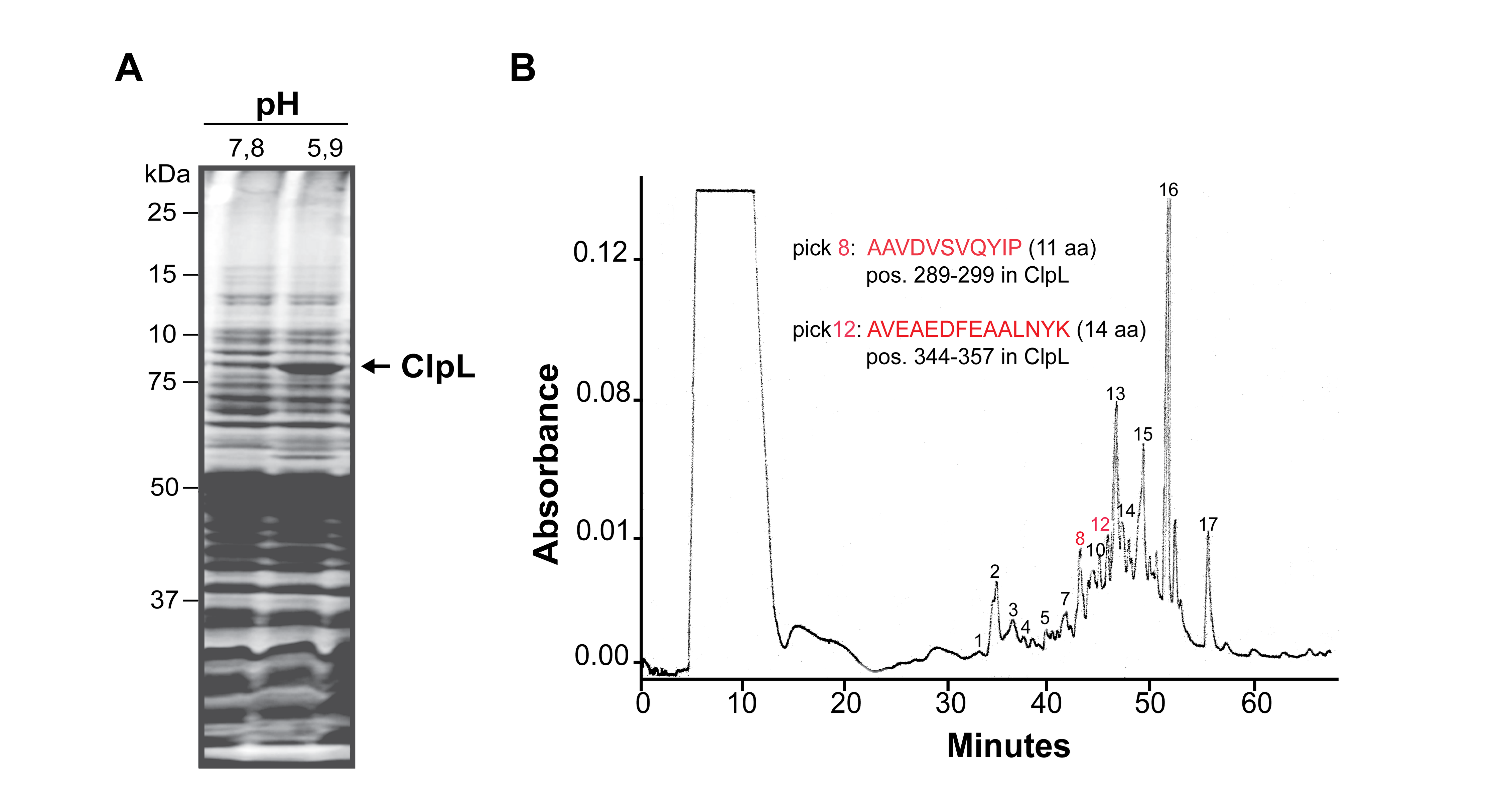

### Fig S7

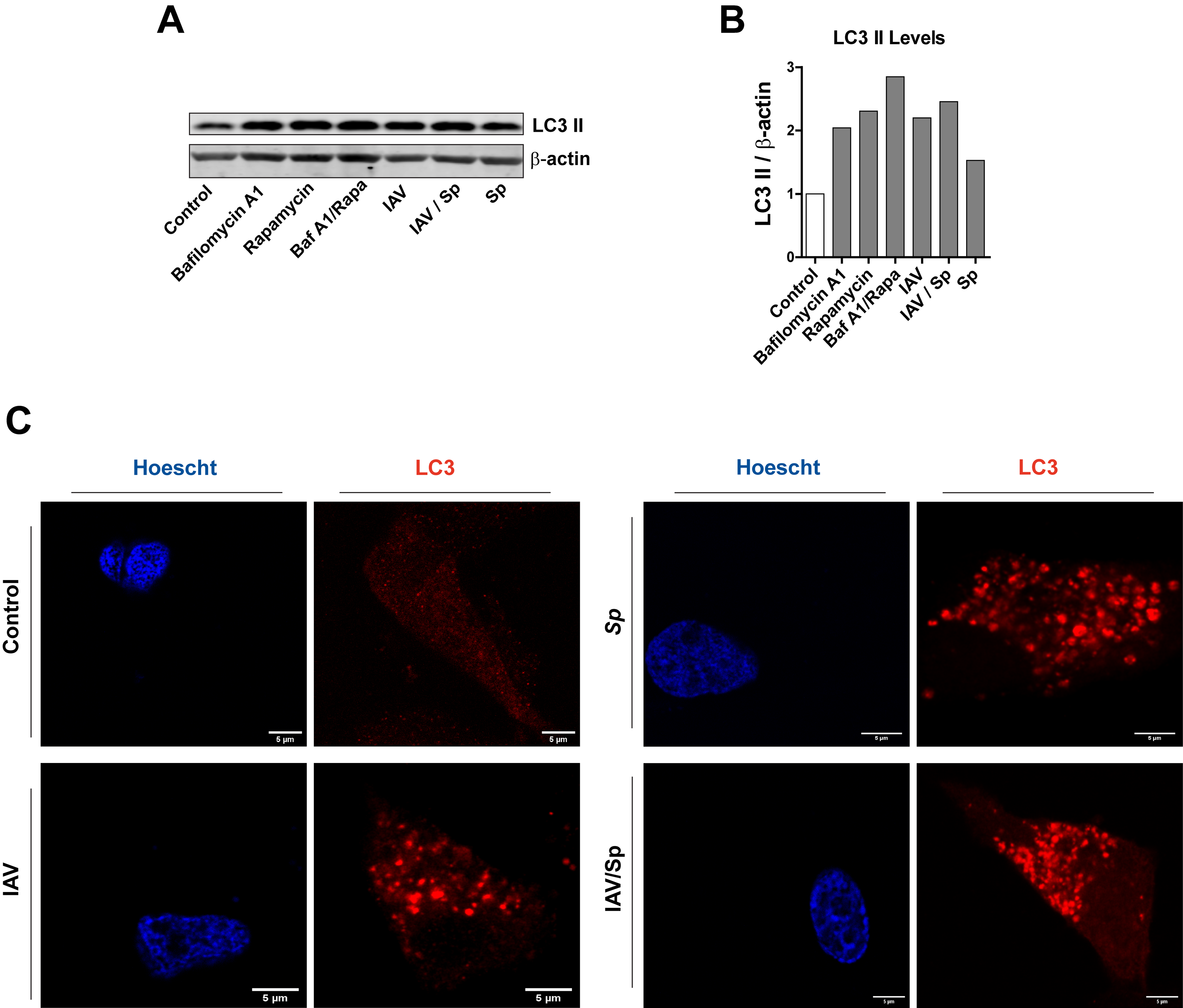
