## Supplementary material for "The pneumococcal two-component system VisRH is linked to enhanced intracellular survival of *Streptococcus pneumoniae* in influenza-infected pneumocytes": Table S1

| **Table S1.** Bacterial strains and plasmids used in this work | | |
| --- | --- | --- |
| ***Strains or plasmids*** | ***Relevant characteristics*** | ***References*** |
| **Strains** |  |  |
| *S. pneumoniae* |  |  |
| R801 | R6 derivative; *hexB* Sm^S^ | 1 |
| R806 | R801, but *rpsL1* obtained by transformation of PCR product amplified by FrpsL and RrpsL primers from the CP1296 chromosomal DNA; Sm^R^ | This work |
| R860 | R806, but ∆*visR::* *kan-rpsL^+^,*  obtained by Janus cassette system (amplified from CP1296 chromosomal DNA) and pneumococcal DNA amplified with the FvisR1, RvisR1, FvisR2 and RvisR2 primers. Km^R^, Sm^S^. | This work |
| R861 | R860, but ∆*visR*, obtained by replacing the Janus cassette with pneumococcal DNA amplified with FvisR1, RvisR1, FvisR2 and RvisR2 primers. Km^S^, Sm^R^. | This work |
| R862 | R806, but ∆*visH::* *kan-rpsL^+^ ,*  obtained by Janus cassette system (amplified from CP1296 chromosomal DNA) and pneumococcal DNA amplified with the FvisH1, RvisH1, FvisH2 and RvisH2 primers. Km^R^, Sm^S^. | This work |
| R863 | 0001, but ∆*visH*, obtained by replacing the Janus cassette with pneumococcal DNA amplified with FvisH1, RvisH1, FvisH2 and RvisH2 primers. Km^S^, Sm^R^. | This work |
| R870 | R806, but ∆*clpL::* *kan-rpsL^+^ ,*  obtained by Janus cassette system (amplified from CP1296 chromosomal DNA) and pneumococcal DNA amplified with the FclpL1, RclpL1, FclpL2 and RclpL2 primers. Km^R^, Sm^S^. | This work |
| R880 | R806, but ∆*psaB::* *kan-rpsL^+^ ,*  obtained by Janus cassette system (amplified from CP1296 chromosomal DNA) and pneumococcal DNA amplified with the FpsaB1, RpsaB1, FpsaB2 and RpsaB2 primers. Km^R^, Sm^S^ | This work |
| R890 | R806, but ∆*sodA::* *kan-rpsL^+^ ,*  obtained by Janus cassette system (amplified from CP1296 chromosomal DNA) and pneumococcal DNA amplified with the FpsaB1, RpsaB1, FpsaB2 and RpsaB2 primers. Km^R^, Sm^S^ | This work |
| RC900 | R801, but *hk01::ery*; Ery^R^. | 2 |
| RC920 | R801, but *hk02:: km*, Km^R^. | 2 |
| RC930 | R801, but *hk03::ery* ; Ery^R^. | 2 |
| RC940 | R801, but *hk04::ery* , Ery^R^. | 2 |
| RC960 | R801, but *hk06::ery*, Ery^R^. | 2 |
| RC970 | R801, but *hk07::ery*; Ery^R^. | 2 |
| RC980 | R801, but *hk08::ery*; Ery^R^. | 2 |
| RC990 | R801, but *hk09::ery* ; Ery^R^. | 2 |
| RC1010 | R801, but *hk10::ery*; Ery^R^. | 2 |
| RC1020 | R801, but *hk11::ery*; Ery^R^. | 2 |
| RC1030 | R801, but *hk13:: ery*; Ery^R^. | 2 |
| RC1040 | R801, but *rr14:: ery*; Ery^R^. | 2 |
| RCM379 | R801 *atpC^A49T^*, Opt^R^ | 5 |
| CP1296 | Rx derivative; *hex mal rpsL1 cbp3::kan-rpsL^+^* (Janus) Km^R^ Sm^S^ | 3 |
|  | D39 but *cpsB::ery,* by transformation with pVA891cpsB , Ery^R^ |  |
| *Escherichia coli* |  |  |
| TOP10 | Amp^R^, Kan^R^, *lacZα*+ selection. F^-^ *mcr*A Δ(*mrr*-*hsd*RMS-*mcr*BC) Φ80 *lac*ZΔM15 Δ*lac*X74 *rec*A1 *ara*Δ 139Δ (*ara*-*leu*)7697 *gal*U *gal*K *rpsL* (Str^R^) *end*A1 *nup*G | Invitrogen |
| DH5α | F -, Φ80dlacZΔM15, Δ(lacZYA-argF) U169, *deoR, recA1, endA1, hsdR17* (rk-,mk+), *phoA, supE44, λ- , thi-1, gyrA96, relA1* |  |
| **Plasmids** |  |  |
| pCR2.1-TOPO | Vector for cloning of PCR products; Ap^R^. | Invitrogen |
| pIRES2-EGFP | Vector for expression of a gene and EGFP on one transcript | Novagen |
| pIRES2-EGFP | Vector for expression of IAV-M2 in eukaryotic cells |  |
| pJDC9 | Integrative vector for *S. pneumoniae;* Ery^R^ | 4 |
| pJDChk01 | pJDC9 containing a 0.3 kb *hk01* amplicon | 2 |
| pJDChk03 | pJDC9 containing a 0.42 kb *hk03* amplicon | 2 |
| pJDChk04 | pJDC9 containing a 0.33 kb *hk04* amplicon | 2 |
| pJDChk06 | pJDC9 containing a 0.32 kb *hk06* amplicon | 2 |
| pJDChk07 | pJDC9 containing a 0.26 kb *hk07* amplicon | 2 |
| pJDChk08 | pJDC9 containing a 0.36 kb *hk08* amplicon | 2 |
| pJDChk09 | pJDC9 containing a 0.41 kb *hk09* amplicon | 2 |
| pJDChk10 | pJDC9 containing a 0.36 kb *hk10* amplicon | 2 |
| pJDChk1)1 | pJDC9 containing a 0.39 kb *hk11* amplicon | 2 |
| pJDChk13 | pJDC9 containing a 0.44 kb *hk13* amplicon | 2 |
| pJDCrr14 | pJDC9 containing a 0.3 kb *rr14* amplicon | 2 |

Abbreviations: Ap^R^: ampicillin resistance; Cm^R^: chloramphenicol resistance; Ery^R^: erithromycin resistance; Km^R^: kanamycin resistance; Str^R^: streptomycin resistance.
