## Supplementary material for "The pneumococcal two-component system VisRH is linked to enhanced intracellular survival of *Streptococcus pneumoniae* in influenza-infected pneumocytes": Table S4

| **Table S4.** Primers used in this work | | |  |
| --- | --- | --- | --- |
| **Primers** | **DNA sequences (5’-3’)** | **Amplified gene** | **Restriction sites** |
| Frpsl | GACGTGCTGACAAATGTTGC | *rpsL* |  |
| RrpsL | AATGTCGACTAGATCTTTCCTTATGCTTTTGGAC | *rpsL* |  |
| F-Janus2 | TTGGATCCGCTAGCCTCGAGAAGCTTGGAACAAGTTATTACTTGAAGATGTCAG | *Janus cassette* | BamHI/NheI/ XhoI/HindIII |
| R-Janus2 | AAGTCGACATCGATAGATCTTCTAGACCCTTTCCTTATGCTTTTGGAC | *Janus cassette* | SalI/ClaI/ BglII/XbaI |
| FvisR1-hlp | CCTCAATCTGCCAACTAAAGG | *spr1475* |  |
| R-MvisR1 | CTTGTTCCGTCGACATCGATCCCAATCTGTTGACGAATGA | *spr1475* | SalI/ClaI |
| F-MvisR2 | GGAAAGGGGTCGACATCGATCAGGGATTGGTAGGATTTATCG | *visH(hk01)* | SalI/ClaI |
| RvisR2-hlp | CCGACCTCAGACTCGATACG | *visH(hk01)* |  |
| F-MvisH1 | TTAATGGTTATCACTGGTGTCAGG | *visR(rr01)* | SalI/ClaI |
| R-MvisH1 | CTTGTTCCGTCGACATCGATAAAGACGACTACGGGAGCG | *visR(rr01)* | SalI/ClaI |
| F-MvisH2 | GGAAAGGGGTCGACATCGATAACGACAGTGCGGATTCAGT | *thrS* |  |
| R-MvisH2 | GAGCTGATGGCGAAGATCAC | *thrS* |  |
| F1M-clpL | CTTGACAGACGGTGTTGACG | *mryA* |  |
| R1M-clpL | GTTGTTGTCGACGGATCCTTACATCAATTGGTTAAATAAATCATCC | *mryA* | SalI/BamHI |
| F2M-clpL | GGAGTTGTCGACGGGATCCGGAAGCAGATATGGAAGATGG | *luxS* | SalI/BamHI |
| R2M-clpL | AATCGATCATCTCGGTAGCC | *luxS* |  |
| FsodA1-Jn | GCCATTGAAGAAAGTCAGAAGTTGAC | *holA* |  |
| RsodA1-Jn | CTGATGTCGACGGATCCAGCGGGTTGTAGTTTCAACAAATGTC | *holA* | SalI/BamHI |
| FsodA2-Jn | ATCTAACTCGAGAGACTCTAATGATAGTTGGAGGGAAGAATTGTTC | *spr0675* |  |
| RsodA2-Jn | TCAAAGGGCTCACCGATTCC | *spr0675* |  |
| FpsaB1-Jn | AGAAAGGGCACGGCTGTAGG | *pepO* |  |
| RpsaB1-Jn | AATCAAAGATCTAAAAAGCTTAACGTATCATAAACTTGTATTCTTCTTGTC | *pepO* | BglII/HindIII |
| FpsaB2-Jn | ATATCAGGATCCAAGCTTATGATTGCAGAATTTATCGATGGATT | *psaC* | BamHI/HindIII |
| RpsaB2-Jn | AACGTCTTCAGGAAGTGGTTCG | *psaC* |  |
| FM2int | CCTATCAGAAACGAATGGGG | *IAV m2* |  |
| FM2-NHE | AATGCTAGCCACCATGAGTCTTCTAACCGAGGTCGAAACGCCTATCAGAAACGAATGGGG | *IAV m2* | NheI |
| RM2-FLAG-ECO | ATAAATGAATTCTTACTTATCATCGTCGTCTTTGTAGTCCTCCAGCTCTATGCTGACAAA | *IAV m2* | EcoRI |
| FgyrA-RT | GGAGATAGTGCTGCCGCTCAACG | *gyrA* |  |
| RgyrA-RT | GGCAAGACCAAGGGTTCCCGTTC | *gyrA* |  |
| FclpL-RT | CGAAACTTGACAGCAGAAGCG | *clpL* |  |
| RclpL-RT | CGGCGTGAGAGGATTTCAGATG | *clpL* |  |
| FvisH-RT | ATTGGTGGAAACGCAGGTCT | *visH* |  |
| RvisH-RT | TAGAGTTCCATCTCACGCGC | *visH* |  |
| FpsaB-RT | GATGAACCCTTTGCTGGGATTG | *psaB* |  |
| RpsaB-RT | TCTTGCTGAGGTCGTGGTGAAC | *psaB* |  |
| *FmurN-RT* | AGGCTGAAACCTTTGGCATTC | *murN* |  |
| RmurN-RT | AGCTTGCTATGAGAAACTCCGC | *murN* |  |
| FaroC-RT | TGTCTTTACTTCGGGCGTTCG | *aroC* |  |
| RaroC-RT | CAGCCATTTCTGGTGGTCCTTA | *aroC* |  |
| FglyA-RT | AACCACATTCAGGAAGCCAAGC | *glyA* |  |
| RglyA-RT | GCTGCCAAATCCATTCCCATAA | *glyA* |  |
